# Genotype and Th2 cells control monocyte to tissue resident macrophage differentiation during nematode infection of the pleural cavity

**DOI:** 10.1101/2021.12.17.472661

**Authors:** Conor M Finlay, James E Parkinson, Brian HK Chan, Jesuthas Ajendra, Alistair Chenery, Anya Morrison, Emma L Houlder, Syed Murtuza Baker, Ben Dickie, Louis Boon, Andrew S MacDonald, Joanne E Konkel, Dominik Rückerl, Judith E Allen

## Abstract

The recent revolution in tissue-resident macrophage biology has resulted largely from murine studies performed in the C57BL/6 strain. Here, we provide a comprehensive analysis of immune cells in the pleural cavity using both C57BL/6 and BALB/c mice. Unlike C57BL/6 mice, naïve tissue-resident Large Cavity Macrophages (LCM) of BALB/c mice failed to fully implement the tissue residency program. Following infection with a pleural-dwelling nematode these pre-existing differences were accentuated with LCM expansion occurring in C57BL/6 but not BALB/c mice. While infection drove monocyte recruitment in both strains, only in C57BL/6 mice were monocytes able to efficiently integrate into the resident pool. Monocyte to macrophage conversion required both T cells and IL-4Rα signalling. Host genetics are therefore a key influence on tissue resident macrophage biology, and during nematode infection Th2 cells control the differentiation pathway of tissue resident macrophages.

**Graphical Abstract:** 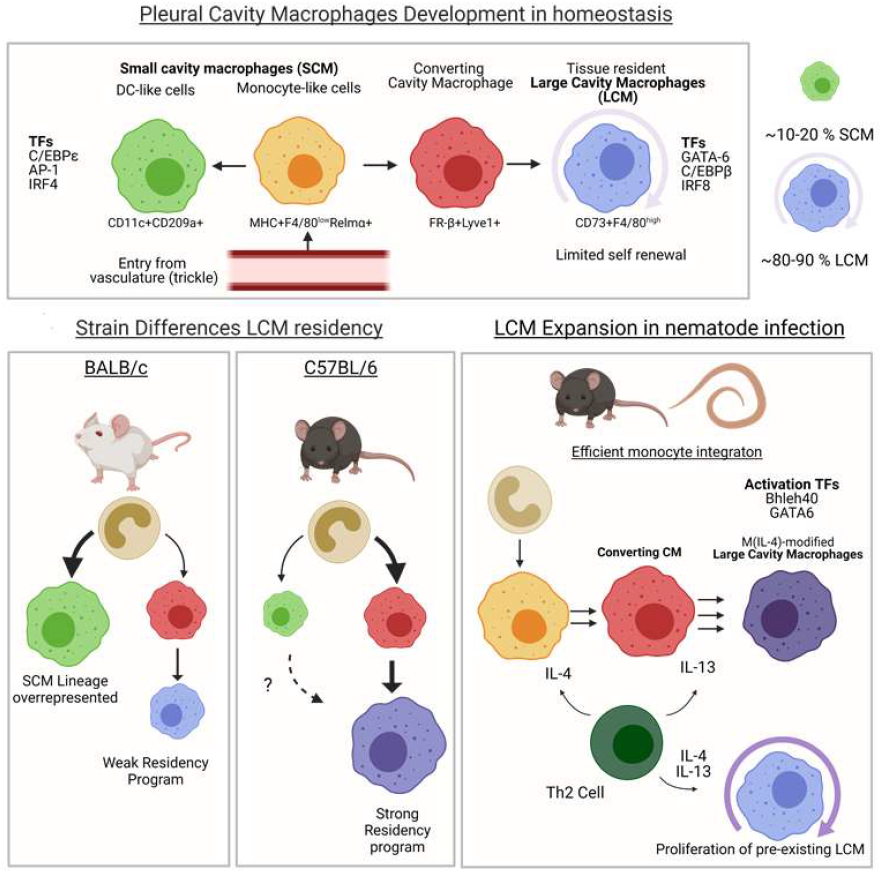

## Introduction

Inflammation of the pleural cavity manifests clinically as pleural effusion, an expansion of pleural serous fluid volume (Miserocchi, 1997), observed in congestive heart failure, pneumonia, cancer, and fibrotic diseases (Feller-Kopman and Light, 2018). Despite major clinical relevance, the immunology of the pleural cavity remains remarkably understudied. Serous fluid is populated predominantly with B cells, T cells and macrophages (MΦ), which subdivide into tissue resident Large Cavity MΦs (LCM), (Buechler et al., 2019; Gautier et al., 2014; Okabe and Medzhitov, 2014) and monocyte-derived Small Cavity MΦs (SCM) (Bou Ghosn et al., 2010; Kim et al., 2016) with evidence of similar heterogeneity in human serous cavities (Chow et al., 2021).

*Litomosoides sigmodontis* is a rodent filarial nematode which infects the pleural space (Finlay and Allen, 2020). Resistance to infection requires IL-4 (Le Goff et al., 2002) and adaptive immunity (Layland et al., 2015), but is host genotype-dependent (Petit et al., 1992). In susceptible BALB/c mice, parasites develop to sexual maturity 50-60 days post infection (p.i) while in resistant C57BL/6 mice infection is established but parasites die before they produce offspring. Using this model we discovered that MΦ cell numbers in the pleural cavity expand in C57BL/6 mice by local IL-4Rα- dependent proliferation without input from the bone marrow (Jenkins et al., 2011). Critically, the same expansion in pleural LCM is not seen to the same degree in infected BALB/c mice, which instead display enhanced monocyte recruitment. Blocking monocyte entry enhances worm killing, linking MΦ phenotypes to disease outcomes (Campbell et al., 2018). Most fundamental insights in MΦ biology over the past decade have been made using C57BL/6 mice. As such, the effect of host genotype on MΦ biology is relatively underexplored. Although genotype has been shown to influence the response of MΦs to Th2 cytokines (Hoeksema et al., 2021; Tang et al., 2020), these studies were not conducted in a natural type 2 immune response setting.

Here, we define MΦ dynamics in the pleural cavity of naïve and nematode infected mice, directly comparing BALB/c and C57BL/6 backgrounds. In naïve mice, we found significant strain differences in LCM but not SCM, with C57BL/6 LCM possessing a stronger resident phenotype. Following nematode infection, monocyte recruitment occurred in both strains, but rapid differentiation into resident LCM was evident only in C57BL/6 mice. Monocyte-derived cells in BALB/c mice were arrested at an intermediate phenotype, unable to activate tissue residency or fully acquire the M(IL-4) activation profile. Unravelling the basis for this strain difference led us to discover that beyond their ability to drive MΦ proliferation (Jenkins et al., 2011), IL-4 and IL-13 produced by T helper cells are required for conversion of recruited monocytes to tissue resident MΦs. This study therefore not only highlights MΦ cell-intrinsic differences between mouse genotypes but reveals an unappreciated role for adaptive immunity in controlling MΦ tissue residency.

## Results

### Nematode infection alters the immune cell profile of the pleural fluid in a strain-specific manner

To better understand the influence of type 2 inflammation on the pleural space we infected resistant C57BL/6 and susceptible BALB/c mice with *L. sigmodontis* (Figure S1A), focusing our analysis on timepoints prior to full parasite clearance in the C57BL/6 strain (Figure S1B, Finlay and Allen, 2020). MRI imaging revealed that infection led to pleural effusion in both strains (Figure 1A and S1C). This was accompanied by an increase in pleural immune cell numbers that was greater in C57BL/6 mice than BALB/c mice (Figure S1D). Mass cytometry of CD45^+^ cells in the pleural lavage showed limited differences in immune cell makeup between the two strains when uninfected but pronounced following infection (Figure 1B). To add statistical power, we performed retrospective analysis of flow cytometric data of pleural immune cells from 360 mice (gating shown in Figure S1E, S1F, and S1G). Neutrophilia was restricted to BALB/c mice and appeared late in infection, while eosinophilia occurred in both strains but was more pronounced in BALB/c mice. B cells were the most abundant pleural immune cells in naïve mice and, while they underwent an initial expansion in both strains, by day 42 in BALB/c mice there was a loss in B cell numbers (Figure 1C, 1D and 1E). CD4^+^ T cells expanded in both infected strains. CD11b^+^ mononuclear phagocytes (MNP, gating shown in Figure S1G) displayed the greatest divergence between the strains, with a higher frequency in C57BL/6 compared with BALB/c in naïve mice. This initial difference was amplified during infection with a much greater expansion of CD11b^+^ MNP in C57BL/6 than BALB/c mice (Figure 1C, 1D and 1E). Notably, mass cytometry revealed pronounced heterogeneity within the MΦ compartment (Figure 1B) between naïve and infected mice but also between the two strains suggesting that CD11b+ MNP undergo phenotypic changes during infection that differ by strain. These results show that nematode infection leads to transformation of the pleural space with major differences in immune cell dynamics, particularly in CD11b^+^ MNP, between resistant and susceptible strains of mice.

**Figure 1.**
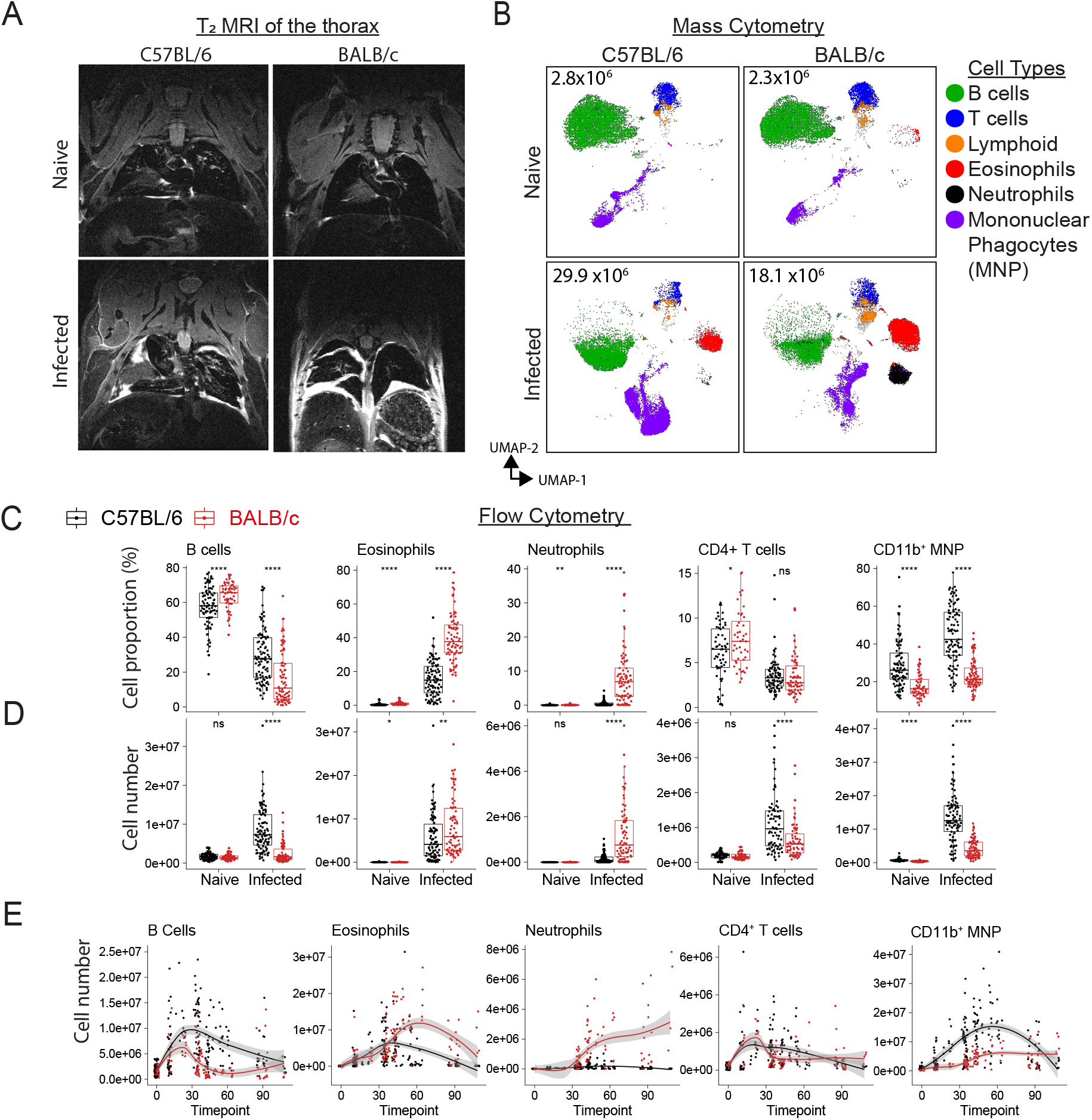
Host genotype dictates cellular immune respon*se to L. sigmodontis* infection in the pleural fluid. Analysis of the pleural fluid of naïve and *L. sigmodontis* infected C57BL/6 and BALB/c mice. A, *T2*-weighted TurboRARE images of the thoracic cavity. B, UMAP of mass cytometric data of CD45^+^ pleural cavity cells, coloured by cell type with mean cell count displayed inside plot. C-D, Summary analysis of pleural cavity flow cytometric data, between day 23-60 p.i.. C, Percentage of cell types as a proportion of total pleural cavity cells D, Data in C adjusted for total cell number. ns = non-significant, **p<0.01, ***p<0.001, ****p<0.0001, t-tests. E, Kinetics of immune cell numbers in infected mice.

### LCM expand in nematode infected C57BL/6 mice but not BALB/c mice

To better capture cellular complexity of pleural CD11b^+^ MNP we used a gating strategy that divided CD11b^+^ MNP into monocytes, SCM, Converting CM and LCM (Figure S1H). Converting CM were a cell type phenotypically intermediate between SCM and LCM (Figure 2A and S2A). SCM made up a greater proportion of CD11b^+^ MNP in BALB/c than C57BL/6 mice, with both strains showing a similar but minor increase in SCM numbers with infection. Monocytes infiltrated to a similar extent in both strains early in infection, but this was accelerated in BALB/c mice after 28 days whilst dropping off in C57BL/6 mice. Converting CM, rare in naïve mice, largely followed the kinetics of monocytes during infection, with a slight delay (Figure 2B, 2C, S2B and S2C). LCMs showed the greatest strain difference, with early and sustained expansion in infected C57BL/6 mice, resulting in a MΦ compartment dominated by LCM (Figure 2B, 2C, S2B). In contrast, there was limited LCM expansion in infected BALB/c mice and, proportionally, a remarkable loss of LCM through infection (Figure 2B, 2C and S2B). The ratio of LCM to recruited MNP populations decreased throughout infection leading to a MΦ compartment dominated by recruited cells in BALB/c mice (Figure 2D and S2D). This ratio negatively correlated with worm recovery (Figure 2E). Together, these data suggest that susceptible BALB/c mice mount a far weaker resident MΦ response to *L. sigmodontis* than resistant C57BL/6 mice. The higher ratio of resident LCM to recruited CD11b^+^ MNP in C57BL/6 than BALB/c mice was also observed in naïve mice (Figure S2D), indicating that strain divergent MΦ responses to infection may accentuate pre-existing differences.

**Figure 2.**
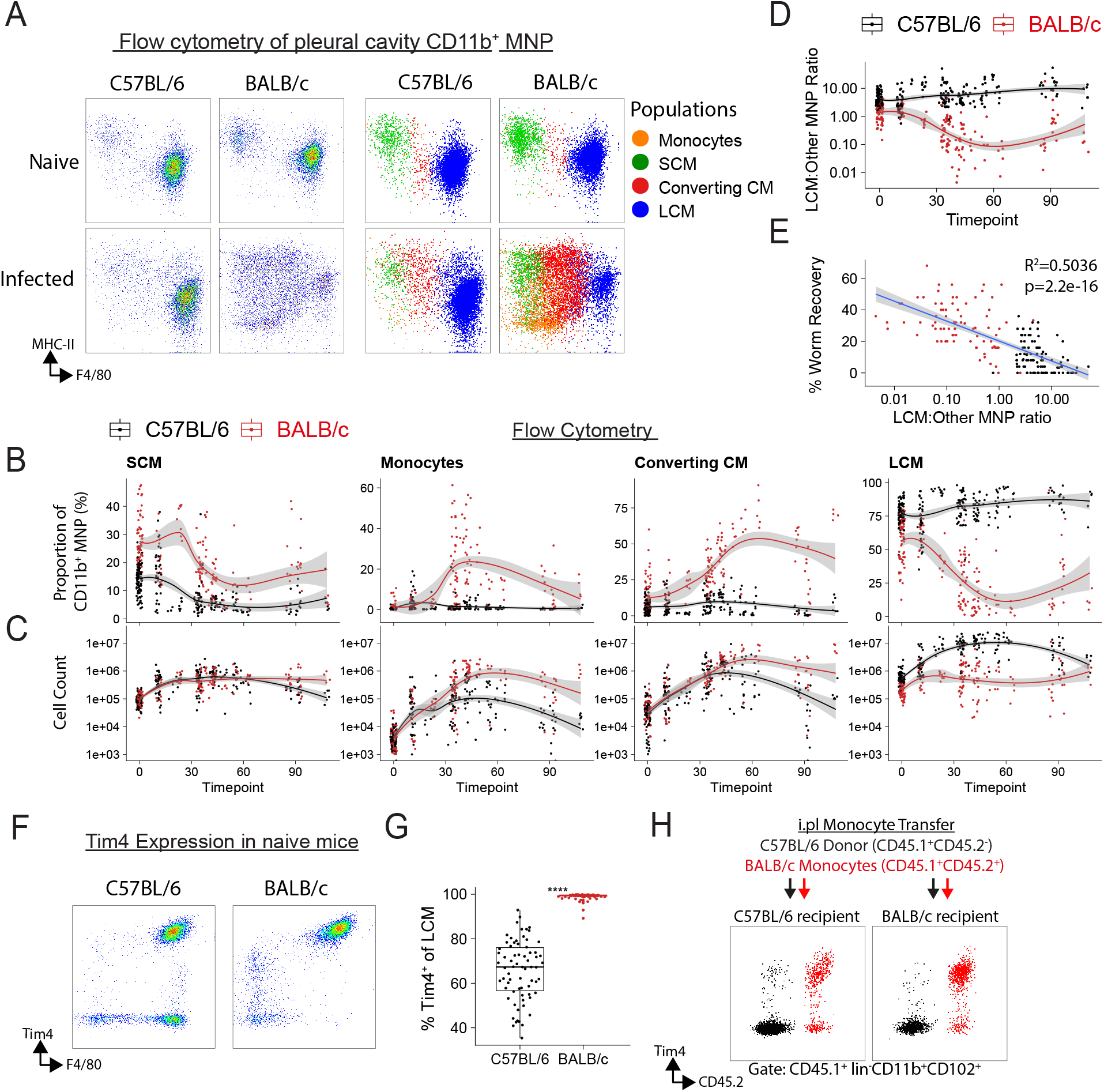
*L. sigmodontis* infection results in LCM expansion in C57BL/6 mice but loss of LCM and increase of monocyte-derived MΦs in BALB/c mice. A-B, Flow cytometric expression of F4/80 and MHC-II gated on Lineage^-^CD11b^+^ MNP, coloured by MNP subpopulation, right. B-C, Kinetics of MNP subpopulations by proportion, C and cell number, D. D, Kinetics of the ratio of LCM to the sum of the other MNP subpopulations. E, linear regression of percentage worm recovery versus the log10 of ratio in D. F-G, Tim4 expression by pleural MΦs in naïve mice. F, representative flow cytometric plot, gated on CD11b^+^Lineage^-^ MNP. G, data summary of Tim4 expression on LCM. H, Expression of Tim4 by CD45.1^+^ LCM following intrapleural transfer of C57BL/6 CD45.1^+^CD45.2^-^ and BALB/c CD45.1^+^CD45.2^+^ bone marrow monocytes into naïve CD45.1^-^CD45.2^+^ C57BL/6 or BALB/c mice.

Expression of Tim4 in naïve mice suggested that pre-existing differences were more than just quantitative. Tim4 was expressed by all BALB/c LCM but a subset of the LCMs in C57BL/6 mice were Tim4^-^(Figure 2F and 2G). Intra-pleural monocyte transfer between the two strains showed that LCM that developed from transferred cells adopted a Tim4 expression profile that matched the donor rather than recipient genotype (Figure 2H). While most BALB/c monocytes had converted to Tim4^+^ LCM, the majority of LCM that arose from transferred C57BL/6 monocytes within this short 4-day development window remained Tim4^-^(Figure 2H). Thus, naïve LCM exhibited cell intrinsic genotype differences.

### Single cell RNA-sequencing of pleural cavity MΦs reveals distinct MΦ populations in naïve mice

C57BL/6 mice had a MΦ compartment that was more ‘resident’ in makeup than BALB/c mice (Figure 3A), and the Tim4 data suggested cell-intrinsic differences. Therefore, to further explore genotype differences, we performed single cell RNA-sequencing (scRNA-seq) on lineage^-^CD11b^+^ pleural CD11b^+^ MNP from naïve C57BL/6 and BALB/c mice (Figure S3A, S3B and S3C). We used single-cell regulatory network inference and clustering (SCENIC) transcription factor (TF) analysis (Aibar et al., 2017), to minimise variance caused by different genetic backgrounds, producing SCENIC TF ‘regulons’ that were used for clustering and dimension reduction (Figure S3D and Figure S4A). This revealed that pleural CD11b^+^ MNP broadly divide into two groups (Figure 3B), which bore similarity to peritoneal LCM and SCM (Figure S4B, Gautiar et al., 2012). LCM had high inferred activity for the TFs CEBP/β, GATA6, MAF/MAFB and various KLF TFs while SCM had high scores for IRF4/5, AP-1 and PU.1 (Figure 3C and 3D). CEBP/β and GATA6 are known to control the residency program of LCM (Cain et al., 2013; Rosas et al., 2014), while IRF4 is needed for development of a subset of SCM (Kim et al., 2016). Relative to SCM, LCM had high expression of the lubricant protein *Prg4*, the B cell chemoattractant *Cxcl13*, various complement/coagulation components (*C1q, Cr3, F5* and *F10*) as well as genes for LCM markers such as *Icam2* (CD102) and *Adgre1* (F4/80)(Figure 3E). Pathway analysis also returned complement and coagulating systems as predicted biological pathways in LCM (Figure S4C), consistent with the role of peritoneal LCMs in coagulation (Zhang et al., 2019). Genes with higher expression in SCM included *Ccr2*, and antigen presentation genes including *Cd74* and MHC genes (Figure 3E). Overall, the gene expression profile was highly consistent with the common designation of LCM as F4/80^hi^MHCII^lo^, and SCM as F4/80^lo^MHCII^hi^.

**Figure 3.**
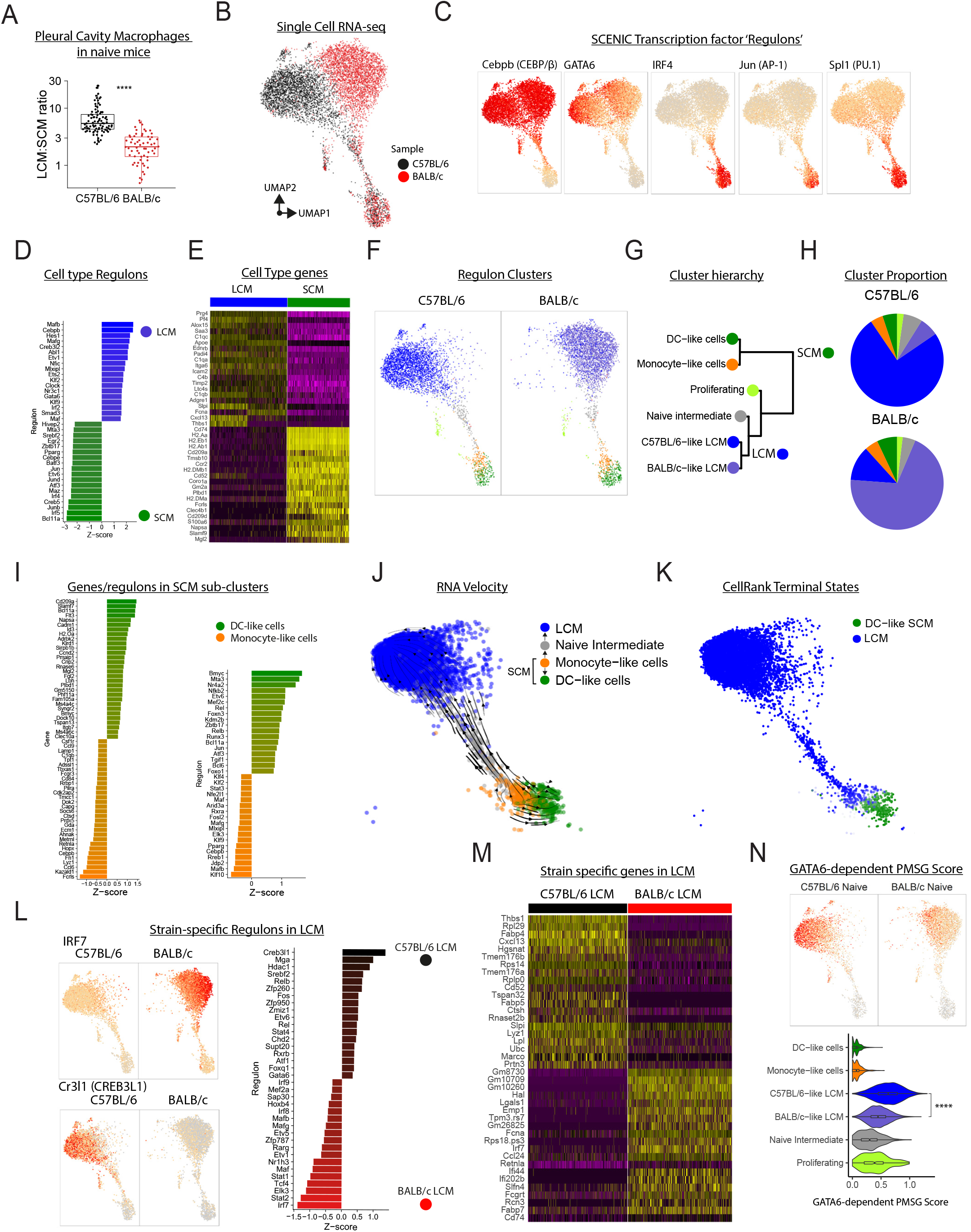
Single cell RNA-sequencing of pleural cavity MΦs reveals novel MΦ populations and that LCM differ by genetic background. A, Ratio of pleural LCM to SCM in naïve mice. ****p<0.0001, t-test. B, UMAP generated from SCENIC TF regulons derived from scRNA-seq data of pleural cavity Lineage^-^CD11b^+^ MNP from naïve mice. C, SCENIC TF regulon activity scores. D, Top differential SCENIC TF regulons between LCM and SCM (both strains). E, Top differential genes between LCM and SCM (both strains). F-H, Hierarchical clustering of SCENIC TF regulons. F, UMAP coloured by cluster, G, cluster dendrogram, and H, cluster proportion. I, Top differential SCENIC TF regulons, left and genes, right between DC-like cells and Monocyte-like cells J-k RNA velocity of cells from C57BL/6 mice. J, vector field stream. K, CellRank predicted terminal states, coloured by likelihood of progression to terminal state. L, Selected SCENIC TF regulon activity scores, left and top strain differential regulons between LCM, right. M, Top differentially expressed genes between C57BL/6 LCM and BALB/c LCM N, GATA6-dependent gene score projected on UMAP, top, and as violin plots separated by cluster, bottom. ****p<0.0001, t-test.

While the LCM-SCM division explained most of the cell heterogeneity, hierarchal clustering separated the pleural MNP compartment into 6 populations. The ‘Proliferating’ cluster, dominated by cell cycle genes was largely excluded from further analysis. SCM-like cells were divided into ‘Monocyte-like cells’ and ‘DC-like cells’ (Figure 3F, 3G and 3H). DC-like cells displayed greater expression of genes associated with DC including *Cd209a* and *Flt3* while Monocyte-like cells had higher expression of core MΦ genes, including *Fcrls* and *Lyz1* (Figure 3I). When purifying cells for scRNA-seq we did not discriminate on CD115 expression, thus the cluster of DC-like cells likely contain CD11b^+^ DC. The Monocyte-like cells had higher activity scores for regulons active in LCM, such as MAF/KLF factors and CEBP/β, albeit to a lower extent than LCM themselves (Figure 3I), suggesting that this population may be destined to develop into an LCM. Fitting this, we identified a rare cluster, ‘Naïve Intermediate’, that sat between Monocyte-like cells and LCMs (Figure 3F). To explore this further, we performed RNA velocity and CellRank fate-mapping analysis to infer trajectory of cell development and predict terminal cell states. This indicated that monocyte-like cells (Figure 3J) develop into either DC-like cells or LCM, which CellRank identified as ‘terminal states’ (Figure 3K).

### More pronounced residency phenotype in naïve C57BL/6 mice than BALB/c mice

SCM clusters from naïve C57BL/6 and BALB/c mice overlapped in UMAP space (Figure 3B), indicating that transcriptionally these cells are largely strain-agnostic. However, LCM populations differentially clustered by strain (Figure 3B and 3G). C57BL/6 LCM had higher regulon activity for CR3L1, FOS, c-REL and STAT4 (Figure 3L) and gene expression for *Thbs1, Fabp4/5, Cxcl13* (Figure 3M). While LCM from BALB/c mice had higher regulon activity for IRF7/9, STAT1/2 and MAF (Figure 3L) and gene expression for *Hal, Lgals1* and *Irf7* (Figure 3M). Pathway analysis showed enrichment for Interferon signalling in BALB/c LCM (Figure S4D). GATA6 controls the residency program of LCM (Rosas et al., 2014), and its regulon activity (Figure 3M) and GATA6-dependent gene score was lower in LCM from BALB/c mice (Figure 3N), suggesting that LCM from BALB/c mice may have a comparatively weak residency program.

### M(IL-4) activation and maintenance of the residency phenotype is greater in nematode infected C57BL/6 mice than BALB/c mice

We expanded our scRNA-seq analysis to include cells from infected animals, which resulted in two new clusters, ‘Converting CM’ and ‘M(IL-4) LCM’ (Figure 4A). Consistent with our flow cytometry analysis (Figure 2), the Converting CM cluster had intermediate gene expression for LCM/SCM markers (Figure 4B) and high expression for *Mrc1* (CD206) and *Lyve1* (Figure 4B and 4C). The M(IL-4) cluster had high expression for LCM markers *Adgre1* and *Icam2* (Figure 4B and 4C), and pathway analysis predicted STAT6 and IL-4 as upstream regulators and oxidative phosphorylation as an upregulated pathway, indicative of M(IL-4) activation (Figure S5A). The proportion of M(IL-4) LCM and Converting CM differed dramatically between strains with M(IL-4) LCM dominating the MNP compartment of infected C57BL/6 mice and Converting CM being the next most abundant (Figure 4D and 4E). CD11b^+^ MNP in infected BALB/c mice were more heterogenous (Figure 4D) with abundant Converting CM and monocyte-like cells, and very few M(IL-4) LCM (Figure 4E).

**Figure 4.**
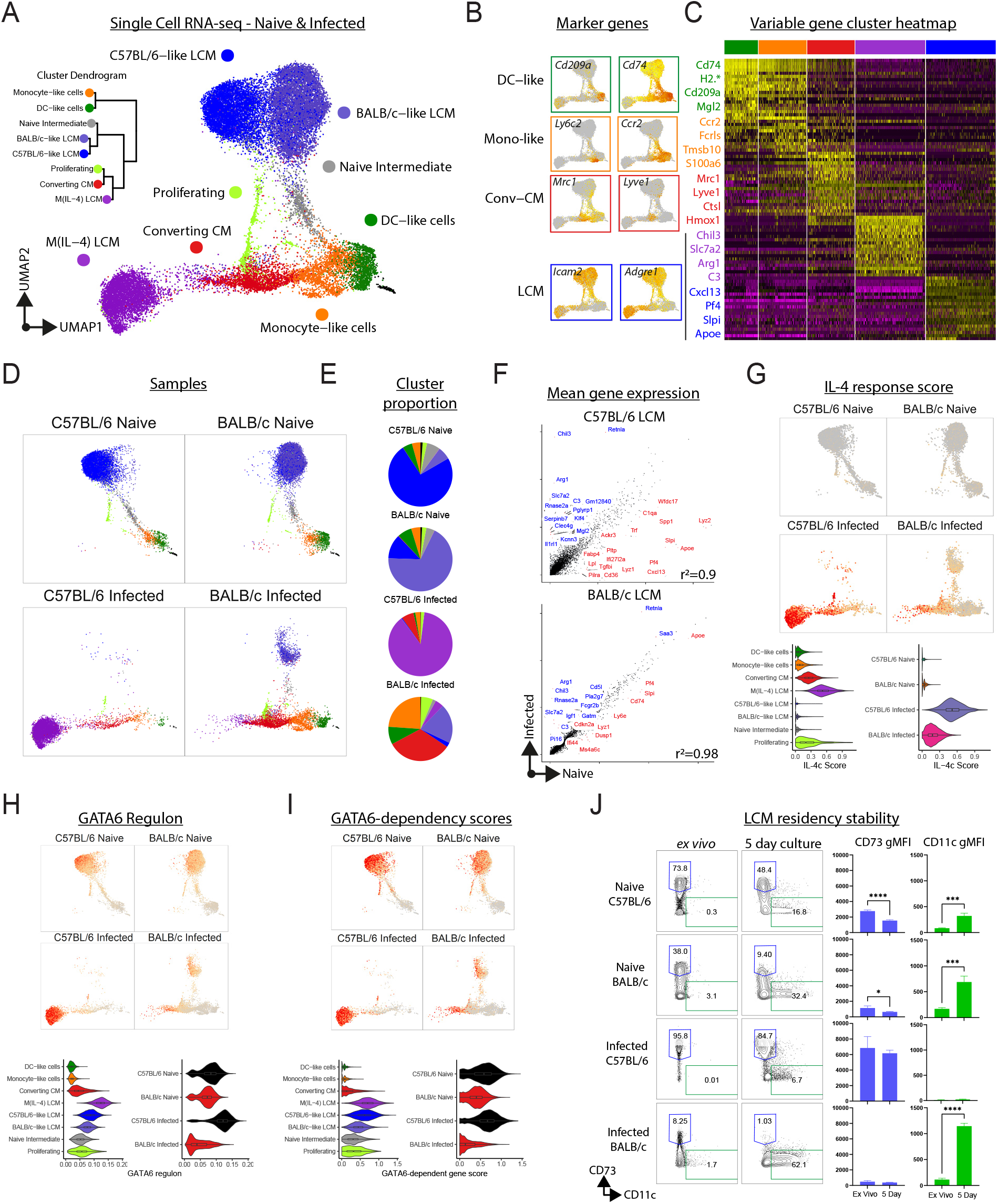
The LCM residency program is stabilised in infected C57BL/6 mice but not in BALB/c mice in which there is an accumulation of immature MΦ. A, UMAP generated from SCENIC TF regulon activity scores from Lineage^-^CD11b^+^ MNP from naïve and *L. sigmodontis*-infected mice, coloured by cluster. Insert, cluster dendrogram. B, Expression of lineage marker genes C, Heatmap of top differential genes in each cluster. Selected genes highlighted to right. D, Data in A, separated by sample. E, Cluster proportion pie charts by sample. F, Scatterplots of mean gene expression by LCM from naïve and infected mice. G, Genelist score for IL-4c response by peritoneal LCM, displayed on UMAP, top or as violin plots separated by cluster or sample, below. H, GATA6 SCENIC TF regulon activity scores on UMAP, above or as violin plots separated by cluster or sample, below. I, Scores for GATA6-dependent genes, on UMAP, above or as violin plots separated by cluster or sample, below. J, Pleural lavage cells were analysed by flow cytometry *ex vivo* and following 5-day *in vitro* culture. Left, expression of CD73 and CD11c gated on CD102^+^ MΦs. Right, gMFI of CD73 and CD11c on CD102^+^ MΦs.

Notably, in infected BALB/c mice, most LCM clustered with LCM from naïve mice rather than with M(IL-4) (Figure 4D) and exhibited less transcriptional change with infection than LCM from C57BL/6 mice (Figure 4F). To test if this reflects a failure by BALB/c LCM to implement the M(IL-4) activation profile we assessed the cells for genes known to be upregulated in LCM following *in vivo* IL-4-complex (IL-4c) delivery (Gundra et al., 2014). M(IL-4) had the highest IL-4c score, followed by Converting CM. Overall, CD11b^+^ MNP from infected BALB/c mice had a more muted IL-4c response (Figure 4G). In C57BL/6 mice, LCM highly upregulated *Chil3* (Ym1), *Retnla* (RELMα) and *Il1rl1* (ST2), while down regulating *Lyz2, Cd36, Pf4* and *Cxcl13* (Figure S5A). These differences were confirmed by RT-qPCR (Figure S5B) and intracellular flow cytometry for Ym1 and RELMα (Figure S5C). Some LCM-associated regulons were not altered by infection (CEBP/β and KLF9), others were lost (RARG and MAF) and still others gained (KLF4, STAT3, FOSB and BHLHE40) (Figure S5A). The loss in activity of cell cycle repressors MAF factors and gain of BHLHE40 may represent a more proliferation permissive state (Jarjour et al., 2019; Soucie et al., 2016), consistent with enhanced LCM proliferation in nematode infected C57BL/6 mice relative to BALB/c mice (Campbell et al., 2018).

We next compared Converting CM to M(IL-4) and found that Converting CM had higher expression of *Mcr1, Lyve1, Cd163* and *Ccl6/9*, with higher relative activity for the SCM-defining regulons IRF4 (Figure S5D). Regulon activity and IPA upstream regulators suggested that interferon signalling was a feature of Converting CM that was not present in M(IL-4) cells (Figure S5D). Interestingly, Gene Ontology scores for response to type 1 Interferons and IFN-γ were higher in CD11b^+^ MNP from BALB/c mice, regardless of infection (Figure S5E).

Converting CM shared some M(IL-4) features but lacked others. To see if this included the LCM residency program, we investigated GATA6. Converting CM had lower GATA6 regulon activity (Figure 4H) and GATA6-dependency scores (Figure 4I) than naïve LCM, but no deficiency in GATA6-independent genes (Figure S5F). Unexpectedly, we found GATA6 regulon activity and GATA6-dependent genes to be enhanced in M(IL-4) LCM over naïve LCM (Figure 4H and 4I). To assess if this reflects an increase in the stability of the LCM residency program we cultured these cells *ex vivo*, removing them from the tissue niche, an approach known to lead to loss in GATA6-dependent residency (Figure S5G, Buechler et al., 2019). Cultured CD102^+^ LCM from naïve mice of both strains lost expression of the GATA6-dependent gene CD73 (Rosas et al., 2014), and gained expression of CD11c, a SCM gene normally silenced in LCM (Bain et al., 2016; Kim et al., 2016). Relative to C57BL/6, BALB/c LCM started with lower CD73 expression and lost this to a greater extent while gaining relatively more CD11c expression (Figure 4J). LCM from infected BALB/c mice, possessed very little CD73 expression to begin with and most of these cells became CD11c^+^ in culture (Figure 3J). LCM from C57BL/6 mice which started with the highest CD73 expression largely retained this expression while not upregulating CD11c (Figure 4J). This result suggests a greater commitment to LCM residency in C57BL/6 mice than BALB/c mice, with infection further stabilising tissue residency in C57BL/6 mice. Taken together these results show that infection of C57BL/6 mice leads to the expansion of M(IL-4) activated LCM, while in BALB/c mice recruited macrophages fail to fully activate the tissue residency and M(IL-4) programs.

### Monocytes integrate into LCM more efficiently in infected C57BL/6 mice than BALB/c mice

We next investigated the hypothesis that Converting CM represent a cell in transition to LCM. RNA velocity and CellRank analysis of the full RNA-seq dataset indicated that monocytes differentiate towards either DC-like cells or the LCM clusters (Figure S6A and S6B), with Converting CM and DC-like cells undergoing the greatest transcriptional change (Figure S6C). Restricting analysis to infected animals also indicated that monocytes transition to LCM (Figure 5A). Computing directional relationships between cell clusters using CellRank and partition-based graph abstraction (PAGA) modelled that monocyte-like cells gave rise to either to DC-like cells or LCM via Converting CM (Figure 5B). CellRank calculated that Monocyte-like cells were the most multipotent cluster with predicted potential (absorption probability) to develop into either DC-like cells or progress down the LCM lineage (Figure 5C), but these probabilities differed by strain. BALB/c Monocyte-like cells had a ∼70% probability of progression to a DC-like cells while C57BL/6-like LCM had an ∼80% probability of moving in the opposite direction towards LCM (Figure 5C and 5D) which reflects the higher proportion of SCM in BALB/c mice. DC-like cells retained a ∼15% probability of progression towards LCM, indicating that this cell may not be a stable terminal state. In contrast, once cells moved into the Converting CM cluster, they appeared largely committed to the LCM lineage (Figure 5C and 5D).

**Figure 5.**
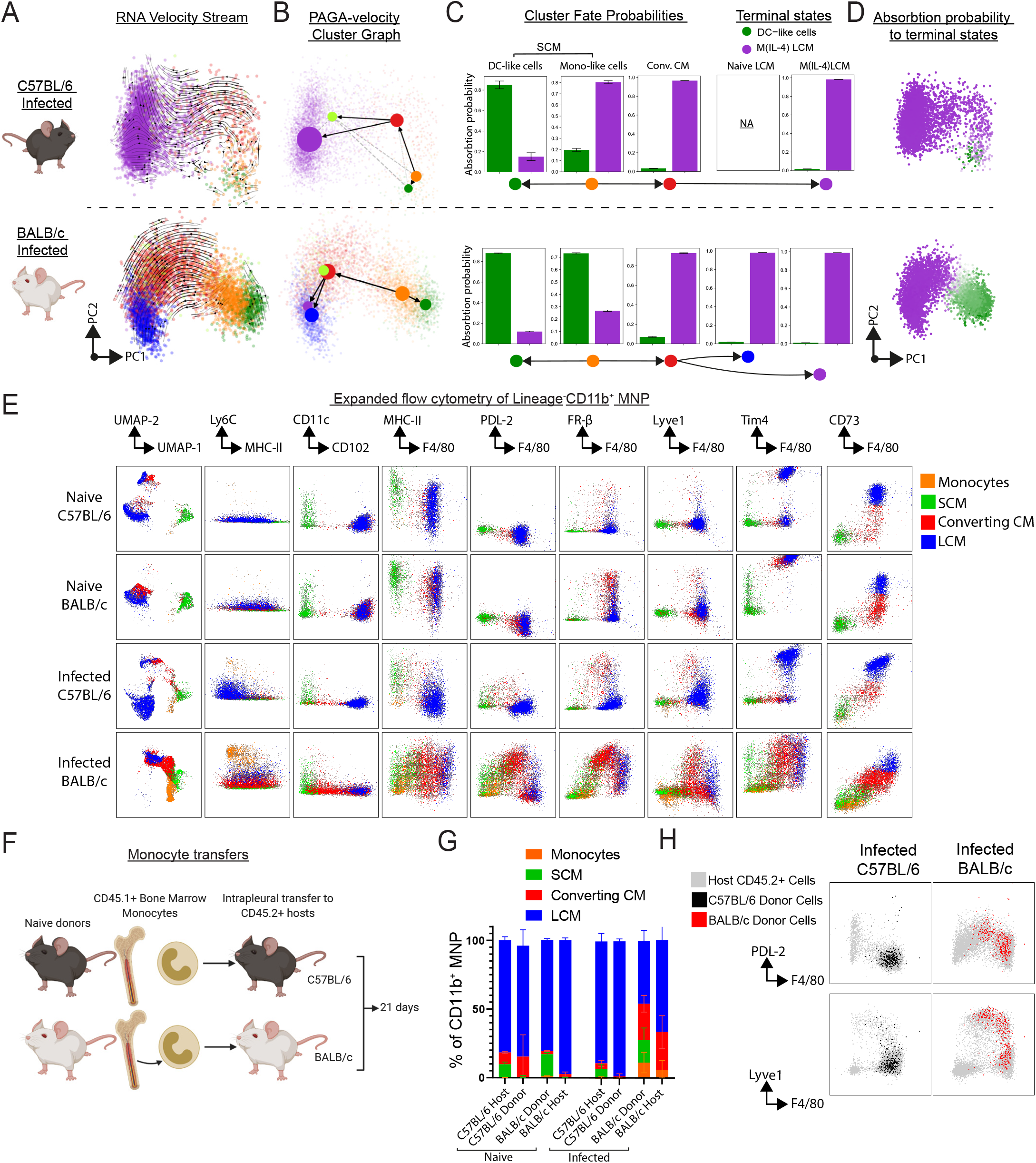
Monocyte differentiation is less efficient in *L. sigmodontis* infected BALB/c mice and they do not acquire residency. A-D, RNA velocity/CellRank analysis. A, RNA velocity vector field stream on variable gene PCA. B, RNA velocity PAGA graphs with clusters as nodes and solid lines depicting high directed edge connectivity. C CellRank fate probabilities for progression to terminal states. D, probabilities displayed on PCA. E, Flow cytometric plots of CD11b^+^ MNP with UMAP generated using 15 parameters. Plots concatenated from 4 naïve and 9 infected mice, coloured by MNP subpopulation. F-H Intrapleural transfer of strain-matched monocytes into naïve and day 14 p.i. mice with recovery of pleural fluid 21 days later. F, Experimental design. G, CD11b^+^ MNP subpopulation proportions for CD45.1^+^ donor and CD45.2^+^ host MΦs. H, Expression of F4/80, PDL-2 and Lyve1 by donor (black/red) and host (grey) MΦs from infected, concatenated from 3 individual mice.

To investigate these possibilities, we first created a flow cytometric panel that captured the spectrum of pleural MΦ states producing UMAP plots like those generated by scRNA-seq (Figure 5E). The panel included markers for DC-like cells (CD11c, CD226), Converting CM (PD-L2, CD206, Folate receptor (FR)-β, Lyve1), LCM commitment (CD102) and LCM terminal maturity (F4/80, Tim4, CD73). Consistent with differences in residency stability seen above (Figure 4J), LCM expressed high amounts of CD73 in C57BL/6 infected mice but CD73^high^ cells were largely absent in infected BALB/c mice (Figure 5E).

Taking these results together, we hypothesized that the efficiency of monocyte integration to LCM differs markedly between C57BL/6 and BALB/c during nematode infection (Figure S6D). To test this, we transferred CD45.1^+^ monocytes into strain matched CD45.2^+^ mice (Figure 5F). Three weeks later all transferred cells had lost their monocyte phenotype (Figure S6E) and none of the transferred cells exhibited a CD11c^+^ DC-like cell phenotype in any recipient (Figure S6F), consistent with the short lifespan of these cells (Bain et al., 2016). Transferred cells were instead committed to the LCM lineage as evidenced by the expression of CD102 (Figure S6G). In naïve mice of both strains and critically in infected C57BL/6 mice the donor cells integrated into the terminal LCM pool (Figure 5G, 5H and S6I). In contrast, donor cells in infected BALB/c mice did not fully integrate and had lower expression of F4/80 and higher expression of the Converting CM markers, Lyve-1 and PDL-2 (Figure 5G, 5H, S6H and S6I), largely phenocopying cells of the recipient they were placed in. These results indicated that in naïve mice and C57BL/6 *L. sigmodontis-*infected mice monocytes can efficiently differentiate into LCM, but this process is less efficient in infected BALB/c mice.

### Genotype of bone marrow-derived cells dictates LCM expansion

LCM tissue residency is imprinted by retinoic acid (RA) produced by serous cavity mesothelial cells that maintains GATA6 expression (Buechler et al., 2019). Thus, alterations in the tissue niche could explain our strain divergent pleural MΦ phenotypes. Although infection raised pleural RA concentrations, this did not differ between the strains (Figure S7A). Nevertheless, to address if MΦ strain differences are immune cell intrinsic or determined by the niche, we transferred bone marrow (BM) between B10.D2 and BALB/c mice (Figure 6A), creating donor matched and mismatched immune systems in myeloablated recipients (Figure S7B). B10.D2 mice share the *L. sigmodontis* resistance phenotype of C57BL/6 mice but the *H*^*2d*^ haplotype of BALB/c mice (Finlay and Allen, 2020). We found the BM genotype (donor), not stromal cell genotype (recipient), to be the major determinant of the increase in LCM numbers following *L. sigmodontis* infection of BM-reconstituted mice (Figure 6B). BM genotype also determined Th2 cell activity in host mice; mice with B10.D2 BM had higher ST2^+^PD-1^+^ and IL-13 expression by CD4^+^ T cells (Figure 6C and 6D). While we did not observe significant differences in worm numbers (Figure S7C), B10.D2 BM reduced parasite fitness (worm length) in susceptible BALB/c host (Figure S7D). In contrast, BALB/c BM did not reverse B10.D2 resistance phenotype, identifying that there are some non-immune determinants for *L. sigmodontis* resistance (Figure S7D). Taken together, these results showed that BM genotype (immune cells) not stromal cells (tissue niche) is the primary determinant of MΦ dynamics during *L. sigmodontis* infection.

**Figure 6.**
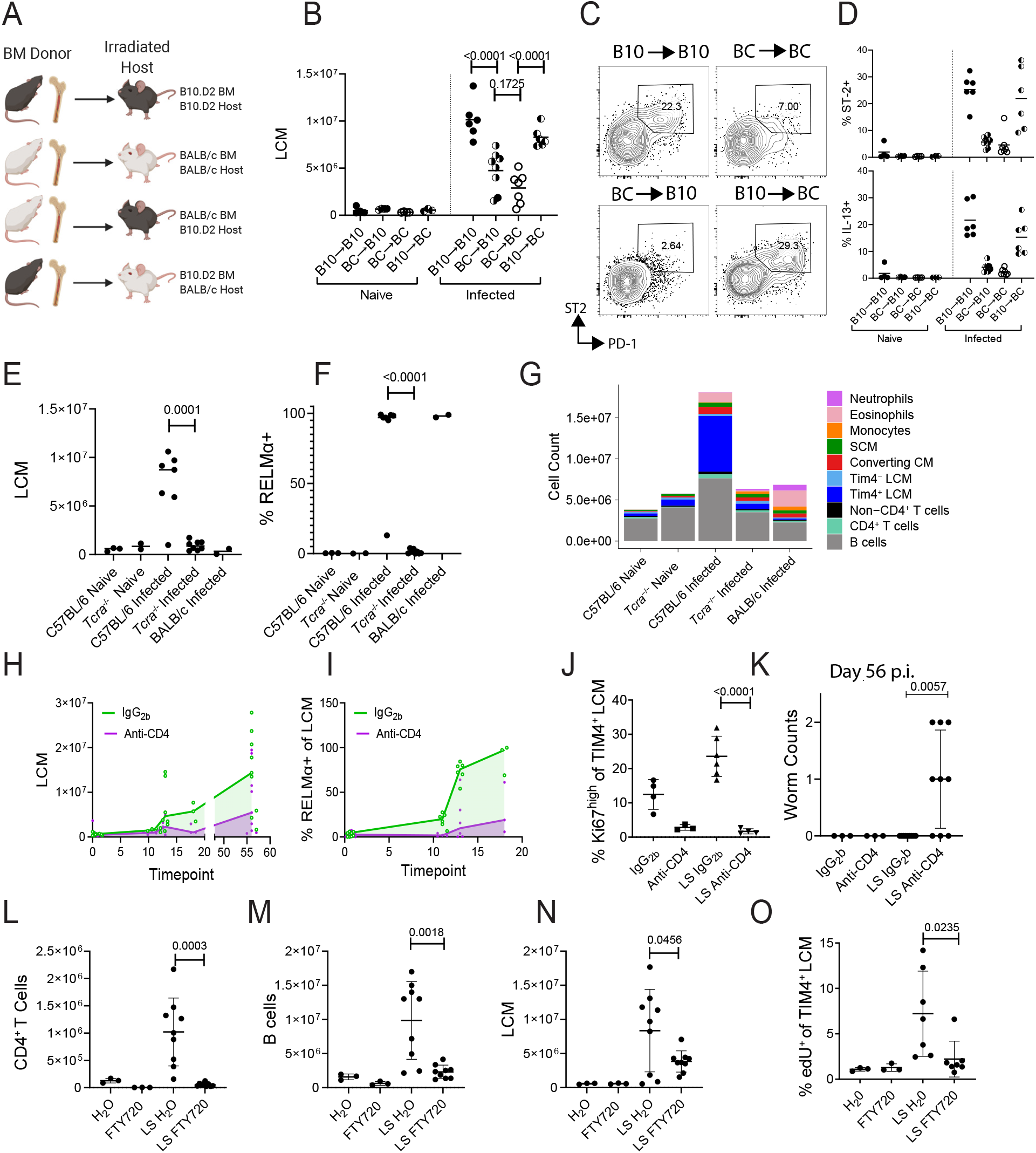
LCM expansion is dictated by the genotype of haematopoietic cells and requires T cells. A-D, Bone marrow transplantation of B10.D2 and BALB/c mice A, Experimental design. B, LCM numbers. C, Flow cytometric plots of PD-1 and ST2 expression by pleural CD4^+^ T cells. D, Expression of ST2 and IL-13 by pleural CD4^+^ T cells. E-G, Pleural immune cells from naïve and day 35 p.i. C57BL/6 and *Tcra*^-/-^ mice. E, LCM numbers, F, percentage RELMα expression by LCM and G, stacked bar charts of total pleural immune cells, coloured in cell type. H-K. Pleural cavity immune cells analysis from C57BL/6 mice given anti-CD4 or isotype control from day 7 p.i.. H, LCM numbers over time, I, percentage RELMα expression by LCM over time. J, Percentage Ki67^high^ of Tim4^+^ LCM taken from naïve and infected mice on day 10 p.i.. K, worm counts on day 56 p.i.. L-O, C57BL/6 mice were treated with FTY720 from day -1 p.i.. L, CD4^+^ T cell numbers, M, B cell numbers, N, LCM numbers in mice on day 33 p.i.. O, Percentage edU incorporation by Tim4^+^ LCM on day 10 p.i.. Horizontal bars represent the median and error bars represent +/- SD. Statistical tests are unpaired t tests, except in A, is a one-way ANOVA with Bonferroni’s correction and K which is a Mann-Whitney U test.

### LCM expansion requires T cells in C57BL/6 mice

Our results demonstrated that *L. sigmodontis*-infected C57BL/6 mice provide a permissive environment for the expansion of the pleural LCM population above homeostatic limits, consistent with previous reports (Jenkins et al., 2013). We next asked what factors in *L. sigmodontis*-resistant C57BL/6 mice facilitate that LCM expansion. Since BM genotype determined LCM expansion (Figure 6B) and this was associated with effector Th2 cell expansion (Figure 6C and 6D), we hypothesized that T cells control MΦ dynamics. To test this, we first used *Tcra*^*-/-*^ mice, which lack mature αβ T cells. These mice were unable to mount LCM expansion during *L. sigmodontis* infection (Figure 6E) and the MΦs present were not M(IL-4) activated (Figure 6F). Indeed, the cellular makeup of the pleural space of infected *Tcra*^*-/-*^ was remarkedly like that of naïve mice, with a limited eosinophilia/neutrophilia (Figure 6G). We next specifically depleted CD4^+^ T cells with antibody and found that this reduced infection-induced LCM expansion (Figure 6H) and LCM RELMα expression (Figure 6I). LCM undergo a proliferative burst around day 10 p.i. in *L. sigmodontis*-infected C57BL/6 mice (Jenkins et al., 2011) and we found that this to be absent in mice depleted of CD4^+^ T cells (Figure 6J). Looking at day 56 in these mice, we found live adult worms in the pleural cavity of infected mice that received anti-CD4 antibodies, whereas the control animals had as expected fully cleared infection (Figure 6K), demonstrating the critical role for T cells in the resistant phenotype of C57BL/6 mice, consistent with recent findings (Wiszniewsky et al., 2021).

The pleural cavity contains resident T and B cells and tertiary lymphoid clusters form within fat deposits in the pleural space (Jackson-Jones et al., 2016). To determine if *de novo* lymphocyte activation in lymph nodes or if local activation of pleural-resident lymphocytes was sufficient for LCM expansion in *L. sigmodontis* infection, we utilised FTY720 which prevents lymphocyte egress from the lymph nodes. Administration of FTY720 reduced pleural CD4^+^ T and B cell expansion in *L. sigmodontis-*infected mice (Figure 6L and 6M). Critically, FTY720 administration also significantly reduced LCM expansion (Figure 6N). Early *L. sigmodontis-*induced LCM proliferation was also reduced by FTY720 (Figure 6O). Collectively these data show that CD4^+^ T cell activation in the lymph nodes followed by entry to the pleural space are required for the expansion and M(IL-4) polarisation of tissue resident LCM in nematode infection.

### Th2 cytokines control monocyte to LCM conversion during nematode infection in C57BL/6 mice

We next assessed the specific role of Th2 cells in two ways, firstly using IL-33 deficiency to prevent Th2 cell expansion (Guo et al., 2015) and secondly using CD11c^+^ cell-specific *Irf4* deficiency to impair Th2 cell development (Williams et al., 2013). We found that Th2 cytokine production (Figure 7A) was significantly reduced in C57BL/6 IL-33LacZGt mice, with no change in IFN-γ (Figure 7B). These IL-33-deficient mice failed to undergo LCM expansion during infection (Figure 7C). The relative proportion of subpopulations that made up the MΦ compartment was similar between IL-33 sufficient and deficient mice despite the overall reduction in CD11b^+^ MNP (Figure 7D). Infection of C57BL/6 *Irf4*^*fl/fl*^ CD11c-Cre^+^ mice resulted in a near total loss of CD4^+^ T cell type 2 cytokine expression, along with enhanced IFN-γ expression, compared to Cre^-^ littermate controls (Figure 7E). Cre^+^ mice also had reduced LCM numbers (Figure 7F) and a loss in RELMα expression by total pleural MΦs (Figure 7G). Cre^+^ *Irf4*^*fl/fl*^ *CD11c-Cre* mice displayed a perturbed MNP compartment, with a greater proportion of monocytes than Cre^-^ animals (Figure 7H) suggesting a failure to transition out of the monocyte state.

**Figure 7.**
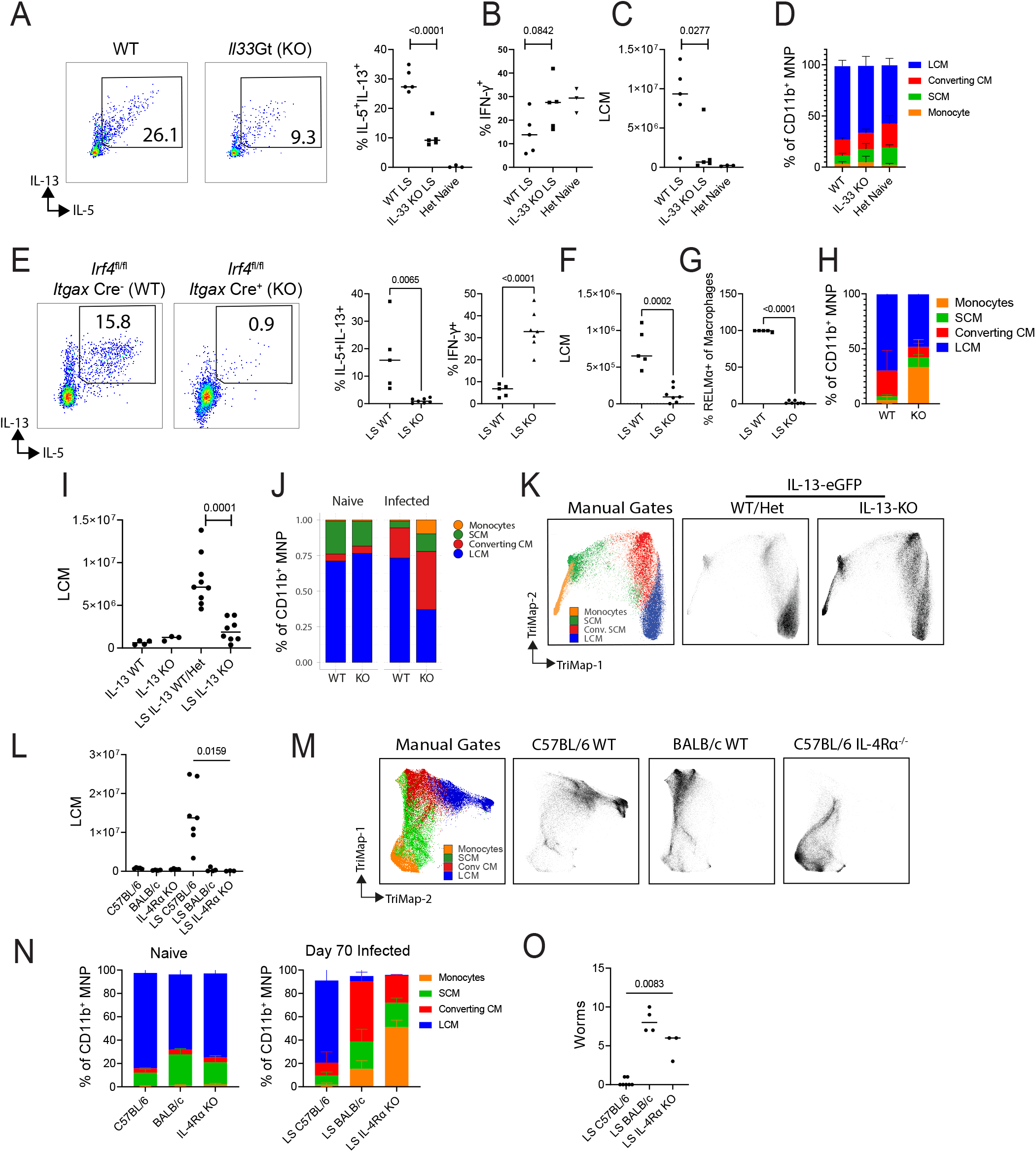
Th2 cells and the cytokines IL-4 and IL-13 control monocyte to LCM differentiation during nematode infection. A-D, Pleural cavity immune cells from IL-33Gt^-/-^ (WT) and IL-33Gt^+/+^ (KO) mice on day 42.p.i., naïve heterozygous mice (Het) were used for visual comparison. A, IL-5 and IL-13 expression by CD4^+^ T cells, data summary, right. B, IFN-γ expression by CD4^+^ T cells. C, LCM numbers and D, MNP subset proportions. E-H, Pleural cavity immune cells from infected Irf4^fl/fl^ *Itgax*-cre^-^ (WT) and Irf4^fl/f^ *Itgax*-cre^+^ (KO) mice on day 41 p.i.. E, IL-5 and IL-13 and IFN-γ expression by CD4^+^ T cells, F, LCM numbers, G, RELMα expression by total MΦs and H, MNP subset proportion. I-K, Pleural cavity immune cells from naïve and infected (LS) IL-13eGFP^-/-^/IL-13eGFP^+/-^ (IL-13 WT/Het) and IL-13eGFP^+/+^ (IL-13 KO) on day 45 p.i.. I, LCM numbers, J, CD11b^+^ MNP subpopulation proportions and K, Trimap of flow cytometric data of CD11b^+^ MNP (concatenated from 3-9 mice per group). L-O, Pleural cavity immune cells from naïve and infected C57BL/6, BALB/c and IL-4Rα^-/-^ mice on day 70 p.i.. L, LCM numbers, M, Trimap of CD11b^+^ MNP (concatenated from 3-8 mice per group), N, CD11b^+^ MNP proportions. O, worm numbers. Horizontal bars represent the median and p values are from unpaired t tests, expect for O, which is a Mann-Whitney U test.

With evidence supporting a role for Th2 cells in LCM expansion in *L. sigmodontis-*infected C57BL/6 mice, we next addressed the specific role of Th2 cytokines. Homozygous IL-13eGFP mice, which lack IL-13, had a reduced expansion of LCMs during infection (Figure 7I) along with a MNP compartment that was dominated by recruited MΦs at the expense of LCM (Figure 7J and 7K), thus making these mice more BALB/c-like. Interestingly infected C57BL/6 mice which lacked IL-13, had a specific loss in LCM but not other immune cells (Figure S7E). This loss suggest that IL-13 plays a specific role in LCM expansion. We next infected C57BL/6 mice which lacked IL-4Rα. Infected *Il4ra*^*-/-*^ mice exhibited a profound loss in LCM (Figure 7L), with a MNP compartment that was dominated by cells at a very early stage of MΦ differentiation, with most cells still falling within the monocyte gate (Figure 7M and 7N). Importantly, the monocyte-dominated phenotype of infected *Il4ra*^*-/-*^ mice did not occur in naïve mice (Figure 7N). Infected *Il4ra*^*-/-*^ mice also still harboured worms at a timepoint in which C57BL/6 mice should have cleared the infection (Figure 7O). These results experimentally validate our previous findings, suggesting that Th2 cells are primary drivers of LCM proliferation and M(IL-4) polarisation (Jenkins et al., 2011) but now reveal a previously unknown role in monocyte to LCM differentiation. We demonstrate that the contribution of IL-4 and IL-13 during type 2 inflammation is even more profound than previously described, and that these cytokines not only drive proliferative expansion of tissue MΦs (Jenkins et al., 2011, 2013) but are required for incoming monocytes to be converted to MΦs.

## Discussion

Our detailed description of pleural cavity MΦs in both naïve and nematode infected mice demonstrate how genetics influence serous MΦ tissue residency. While our findings largely agree with the division of serous cavity MΦ into SCM (monocyte-derived) vs LCM (tissue resident) they reveal a differentiation pathway from monocytes to LCM that differs between mouse strains. The transcriptional similarity between C57BL/6 and BALB/c SCM suggested that the core MΦ program (controlled by PU.1, C/EBPs and AP-1 binding) is relatively stable between these strains consistent with limited differences between C57BL/6 and BALB/c MΦs in response to TLR4 agonists (Link et al., 2018) and IL-4 (Hoeksema et al., 2021). However, in addition to the core MΦ program, the enhancer landscape can define a tissue-specific residency program (Gosselin et al., 2014). Here we show that the residency program differed significantly by strain with LCM from C57BL/6 mice having a ‘stronger’ residency phenotype than BALB/c mice. Nematode infection exaggerated these strain differences in LCM residency, with MΦs in C57BL/6 mice further increasing activity for residency TFs including GATA6 (Buechler et al., 2019), Bhlhe40 (proliferation (Jarjour et al., 2019)) and KLF4 (immunological silencing (Roberts et al., 2017)). In contrast, BALB/c mice displayed a near complete loss in residency following infection. Bone marrow transplantation demonstrated that this divergence in residency phenotype between strains was determined by immune cell genotype, excluding a direct role for the tissue niche in these strain differences, at least during infection.

Counter to the traditional view of the type 2-prone BALB/c mouse, during *L. sigmodontis* infection the Th2 response is actively suppressed in this strain resulting in a mixed Th1/hyporesponsive Th2 response (Knipper et al., 2019). While not the focus of this study, our findings help explain the complex susceptibility phenotype of BALB/c mice (Finlay and Allen, 2020). The failure of these mice to develop fully resident tissue MΦ, leads to an accumulation of PD-L2^+^ MΦ and blockade of their entry enhances worm killing (Campbell et al., 2018). These PD-L2^+^ cells will promote the PD-1-dependent hypo-responsive T-cell phenotype found in both human and mice believed to be largely responsible for failure to kill the parasites (Babu et al., 2006; Knipper et al., 2019; van der Werf et al., 2013). In contrast, we find that the potent Th2 cell response in C57BL/6 mice is responsible for parasite killing consistent with the requirement for IL-4 and T cells in host resistance (Le Goff et al., 2002; Wiszniewsky et al., 2021).

A major unexpected finding of our study was that Th2 cells promoted the adoption of tissue residency by monocytes. Our previous finding that IL-4 and IL-13 drive the proliferative expansion of tissue resident MΦs was made in *L. sigmodontis*-infected C57BL/6 mice (Jenkins et al., 2011) and although we appreciated that some of the proliferating cells had bone marrow origins (Campbell et al., 2018) we failed to observe monocyte recruitment during infection. Our finding here suggest that in the highly polarised Th2 environment of the nematode-infected C57BL/6 pleural cavity, monocytes integrate so rapidly into the LCM pool that they are only seen as rare cells by flow cytometry. Although LCM expansion via proliferation is a feature of many type 2 immune responses (Rückerl and Allen, 2014) monocyte integration, not proliferation can account for increased numbers of M(IL-4) MΦs, as seen in the helminth-infected liver (Nascimento et al., 2014). We propose a model of coexisting proliferation and monocyte integration, whereby the size of the resident MΦ niche is controlled primarily by growth factor availability, acting on both pre-existing resident MΦs to induce proliferation, and monocytes to promote residency integration. In the context of type 2 immunity IL-4Rα, not CSF-1R, signalling, act as primary growth stimuli for LCM expansion (Jenkins et al., 2013). In this model, only in the most type

2 polarised environments (i.e., a strong adaptive Th2 response) is the myeloid compartment dominated by expansion of resident MΦs. In this setting, Th2 cells drive not only proliferation of the existing population but conversion of monocytes to MΦ along with enhancement of the tissue residency program. When Th2 cells, or IL-4Rα signalling is insufficient, as we observed in BALB/c mice or type 2 impaired C57BL/6 mice, monocytes or monocyte-derived MΦ accumulate at earlier stages of maturity. Loss of IL-4Rα signalling had no effect on homeostatic LCM maintenance, indicating that a function of Th2 cytokines during type 2 immune responses is to remove homeostatic constraints that limit resident MΦ numbers. The fluid phase nature of the serous cavities may further facilitate this expansion as it lacks the structural features of other tissues proposed to limit MΦ numbers (Guilliams et al., 2020).

*L. sigmodontis* infection, regardless of host genetic background, elicits inflammatory monocyte recruitment but the dominant Th2 cell response in C5BL/6 mice favours tissue resident MΦ, while recruited MΦ phenotypes dominate in infected BALB/c mice. The later resembles type 1 peritonitis models, with the loss of resident MΦs (MΦ disappearance reaction) in favour of inflammatory monocyte recruitment (Zhang et al., 2019). Converting CM also resemble thioglycolate-elicited inflammatory MΦs that are subsequently given IL-4c (Gundra et al., 2017). The MΦ disappearance reaction may be an evolutionary adaptation to remove proliferative resident MΦs that promote growth of microorganisms (Huang et al., 2018; Wang et al., 2020). The presence of type 1 inflammation in *L sigmodontis* infected BALB/c mice may explain the loss of residency as mice with enhanced interferon-γ signalling have a profound loss in LCM tissue residency (Wang et al., 2020). Further, interferons including IFN-γ are generally anti-proliferative (Gajewski et al., 1988; Wu et al., 2018). These findings are consistent with the reversal of nematode-induced peritoneal LCM expansion by co-infection with *Salmonella* (Rückerl et al., 2017), perhaps reflecting the evolutionary need for type 1 immunity to take precedence over type 2 responses because of the greater threat posed by bacteria over helminths (Graham et al., 2005). Indeed, Th2 cell-dependent monocyte to macrophage conversion may only be clearly evident in settings like the pleural cavity of *L. sigmodontis* infected C57BL/6 mice, with limited pro-inflammatory signals to block integration. The type 1 – type 2 (or M1/M2) dichotomy could thus be expanded to include that anti-microbial M1 MΦ are refractory to proliferation and fail to integrate into the resident pool, while M(IL-4) MΦ integrate into the resident pool and if needed expand through proliferation to fulfil a functional role such as tissue repair or parasite control. This dichotomy is supported by the uncoupling of macrophage inflammation from self-renewal described for alveolar macrophages (Zhu et al., 2021). Finally, our data suggest that innate tissue-specific niche signals are only part of the story in monocyte conversion to tissue-resident MΦ and that adaptive immune cells can take control of these pathways.

## Author contributions

C.M.F. and J.E.A. conceptualized and supervised the study and wrote the manuscript. C.M.F. and J.P. conducted and analysed most experiments. C.M.F. conducted computational analysis. J.P. provided computational analysis. D.R. provided conceptual and technical assistance. B.H.K.C maintained the parasite lifecycle. J. P., B.H.K.C, J.A., A.C, A.M. and J.K. provided technical assistance. B.D. designed the MRI protocol. L.B., E.H. and A.S.M. provided key resources. S.M.B. provided computational assistance. All authors edited and approved the manuscript.

## Acknowledgments

We thank Qiang Shi for sharing code, Mudassar Iqbal for advice on transcription factors, Pete Cook for advice on mass cytometry, Matthew Hepworth, Stephen Jenkins and P’ng Loke for constructive scientific discussion. William Agace, Kara Filbey and John Grainger provided mouse lines. This work was funded Medical Research Council-UK (MR/K01207X/2,) and Wellcome Trust (106898/A/15/Z).

## Conflict of interest

The authors declare no competing interests

---

**Figure S1.**
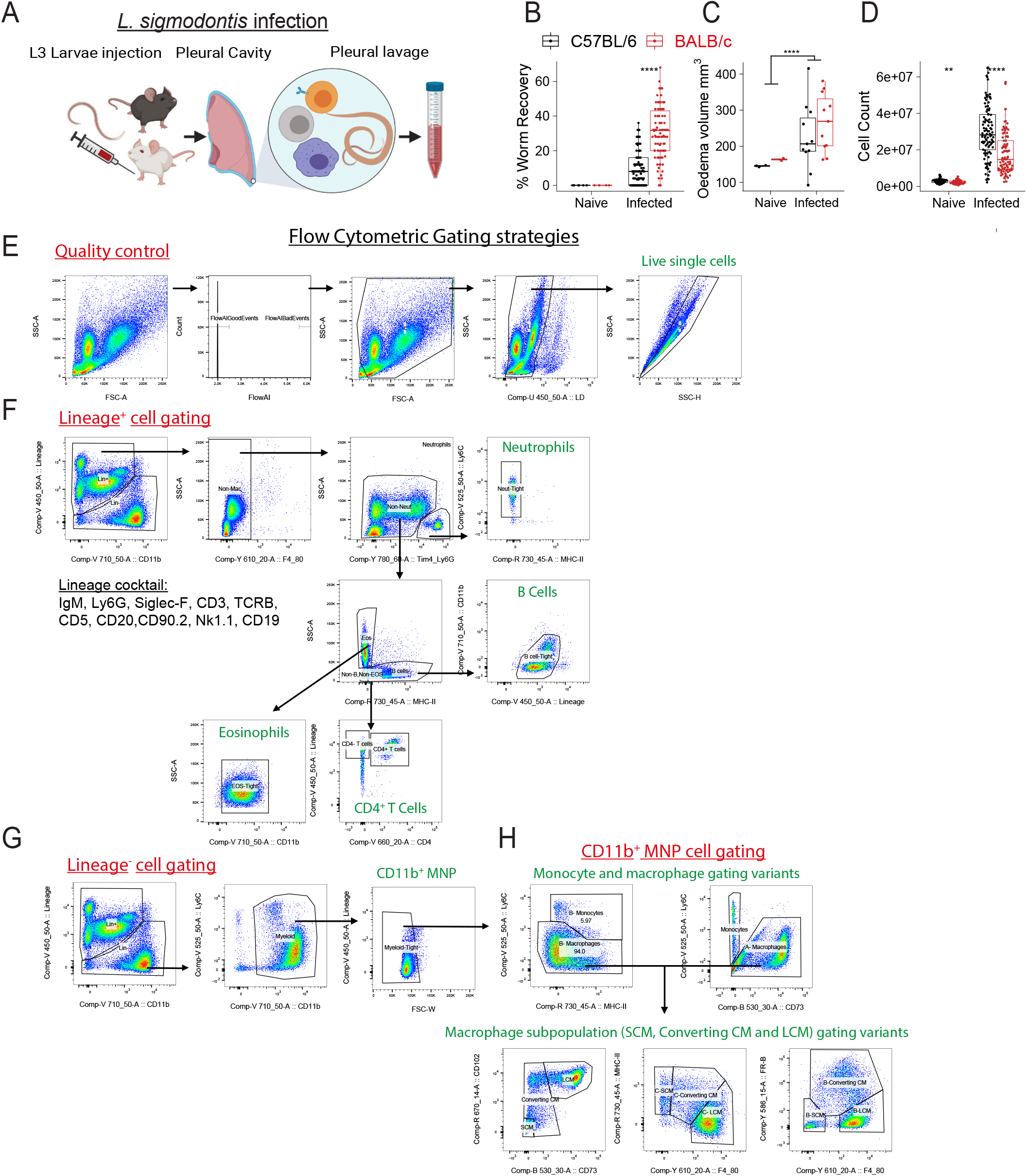
Analysis of pleural cavity immune cells from naïve and *L. sigmodontis* infected C57BL/6 and BALB/c mice. Related to Figure 1. A, Schematic of *L. sigmodontis* infection and recovery of pleural fluid. B, Worm recovery as a percentage of infection dose in C57BL/6 mice and BALB/c mice between day 23-60 p.i. ****p<0.0001, Kruksal-Wallis test. C, Oedema volume calculated from MRI images. ****p<0.0001, t-test grouped by infection status. D, Pleural lavage cell count from naïve and infected C57BL/6 mice and BALB/c mice. **p<0.01, ****p<0.0001, t-tests. E, Quality control gates for selection of debris-free live single cells. For some experiments, cells with staining artefacts were excluded using the FlowAI Flowjo plugin. F, Gating strategy for lineage^+^ cells G, Gating strategy for lineage^-^ cells H, Gating strategies used to identify monocytes, SCM, Converting CM and LCM using different sets of markers that were used in individual experiments. Each gating variants produced similar subpopulation proportions between experiments.

**Figure S2.**
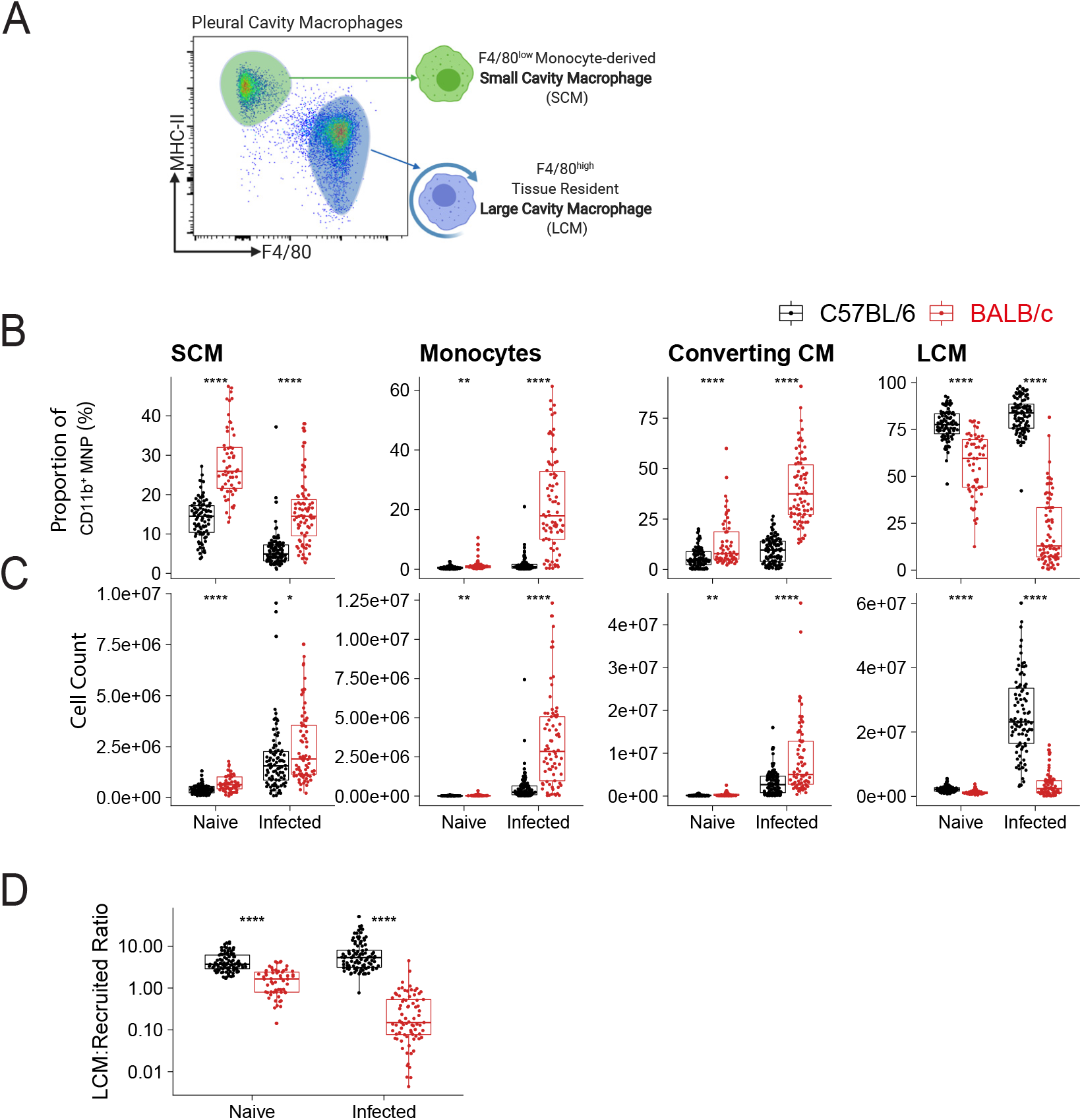
MNP subpopulations in naïve and infected C57BL/6 mice and BALB/c mice. Related to Figure 2. A, Major MΦ subsets in the pleural space, gated on live, single, lineage^-^, CD11b^+^ MNP. B-D, Summary analysis of flow cytometric data of MNP subpopulations in naïve and infected C57BL/6 mice and BALB/c mice, between day 23-60 p.i.. B, Percentage of indicated cell types as a proportion of single, live lineage^-^CD11b^+^ MNP and C, total cell number of individual MNP subpopulations *p<0.05, **p<0.01,****p<0.0001, t-tests. D, ratio of LCM to the sum of the other subpopulations (monocytes, SCM and converting CM, ‘Recruited’), ****p<0.0001, t-test.

**Figure S3.**
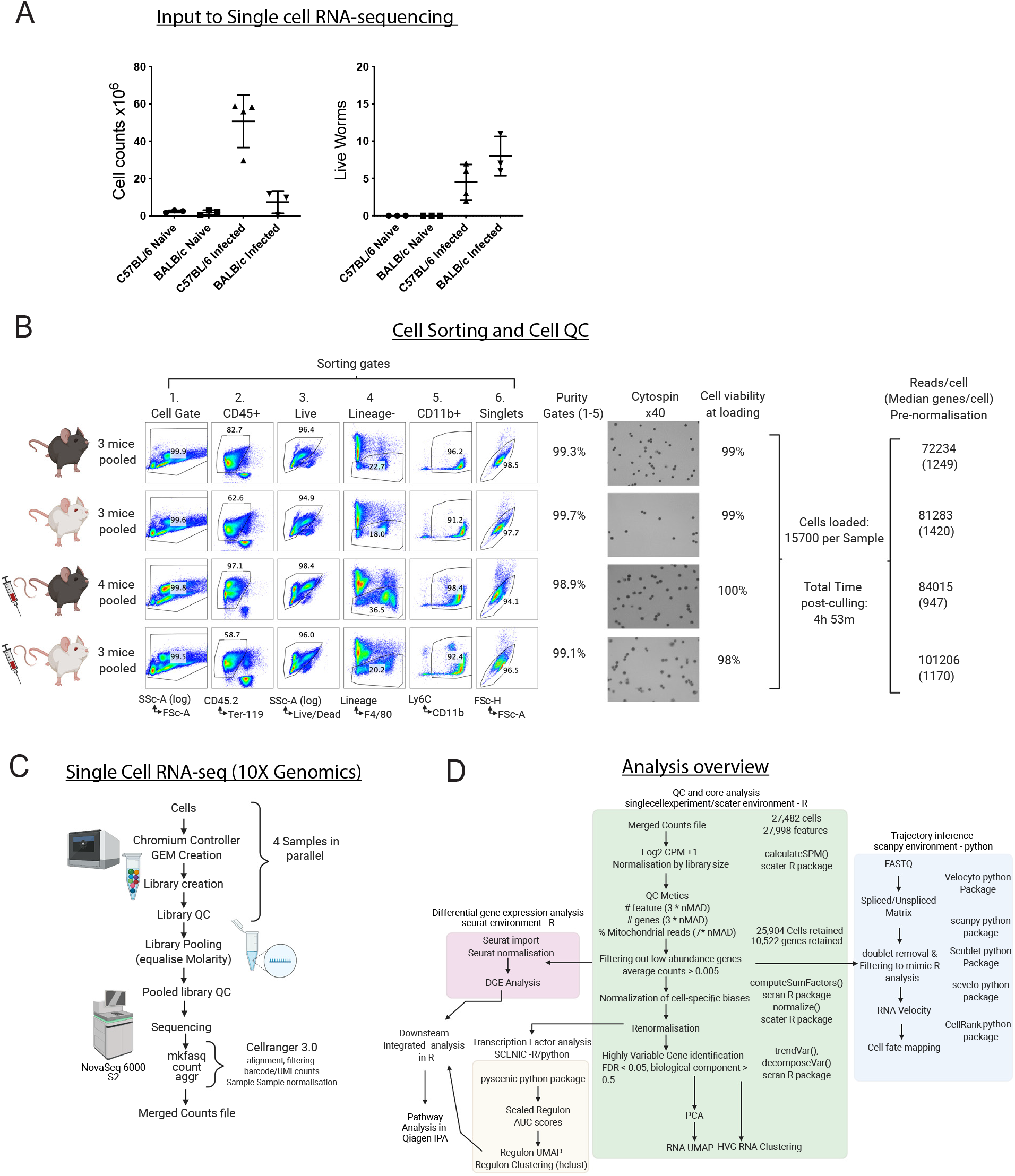
Experimental design and analysis overview of single cell RNA-sequencing experiment. Related to Figure 3 and 4. A, Cell counts and parasite numbers of mice used as source of cells for single cell RNA-seq experiment prior to equalised pooling. B, Overview of cell sorting and quality control for cells used as an input to single cell RNA-seq experiment. C, Overview of 10X genomics single cell RNA sequencing pipeline. D, Overview of quality control, cell/gene filtering and normalisation of initial counts file prior to downstream analysis.

**Figure S4.**
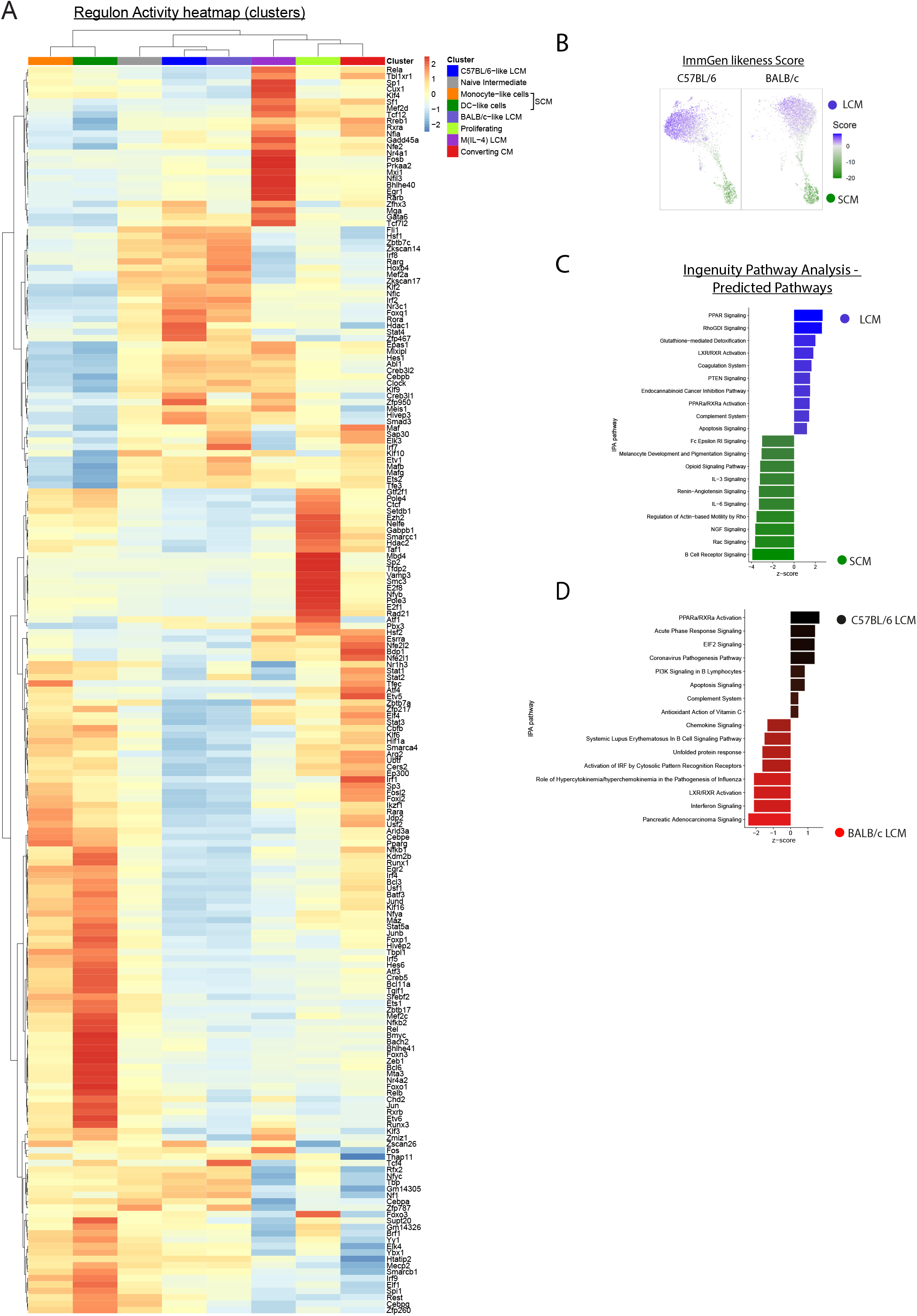
Scenic TF regulon activity and pathway analysis generated from single cell RNA-seq dataset. Related to Figure 3 and 4. A, Heatmap of mean regulon activity scores in each cluster for all 195 regulons (Ward’s method), scaled by row (regulon). B, Weighted similarity score of pleural MΦs for peritoneal LCM (positive values, blue) and SCM (negative values, green) from ImmGen microarray dataset V1. C, Ingenuity pathway analysis displaying significant differential predicted pathways in LCM and SCM. D, Ingenuity pathway analysis displaying significant differential predicted pathways in C57BL/6 LCM and BALB/c LCM.

**Figure S5.**
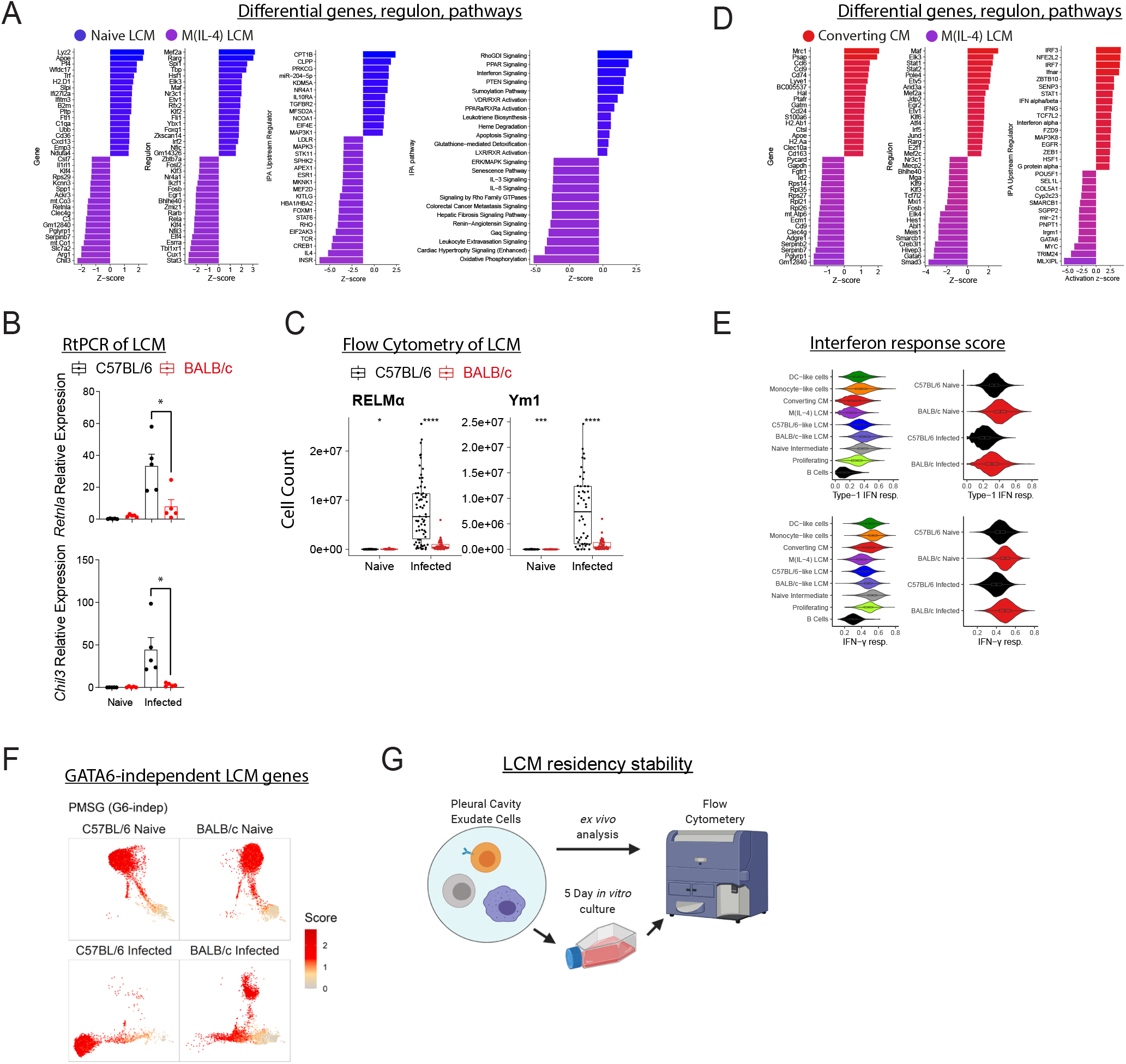
Related to Figure 4. A, Comparison of Naïve LCM and M(IL-4) clusters, with top differentially expressed genes (adjusted p<0.001), left, regulons (adjusted p<0.001), middle, IPA pathway analysis upstream regulators (p value of overlap<0.01), right. B, Quantitative-PCR for *Retnla* and *Chil3* by FACS-sorted LCM from naïve and infected (Day 35 p.i.) C57BL/6 and BALB/c mice, relative to naïve C57BL/6 LCM group. C, RELMα^+^ and Ym1^+^ LCM from naïve and infected C57BL/6 and BALB/c mice, by intracellular flow cytometry. D, Comparison of converting SCM and M(IL-4) clusters, with top differentially expressed genes (adjusted p<0.001), left, regulons (adjusted p<0.001), middle, or IPA pathway analysis predicted pathways, right. E, Gene expression scores for Gene Ontology terms ‘cellular responses to type 1 interferons’ (GO:0071346) and ‘response to interferon-gamma’ (GO:0034341) by cells in the RNA-seq dataset. Data presented as violin plots, separated by cluster, left and sample, right. F, Gene scores for GATA6 independent LCM specific genes. Scores are displayed on the regulon UMAP embedding. G, Experimental design schematic: Pleural lavage cells from naïve and infected (n=3/group) C57BL/6 or BALB/c mice were analysed by flow cytometry *ex vivo* and following 5-day *in vitro* culture.

**Figure S6.**
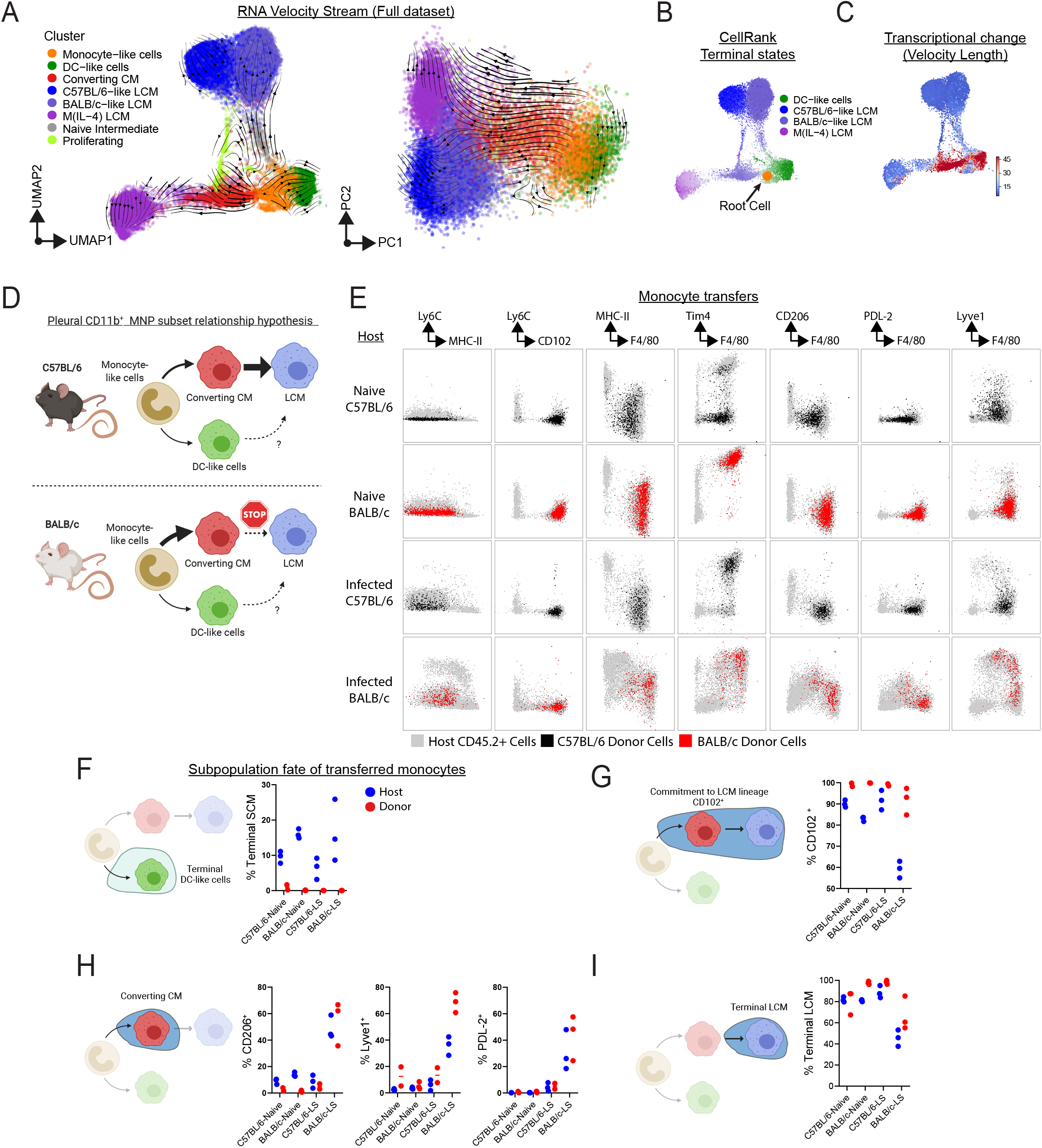
Related to Figure 5. A-D, RNA velocity and CellRank analysis of full RNA-seq dataset. A, RNA velocity vector field stream projected on SCENIC regulon UMAP, or variable gene PCA. B, CellRank predicted terminal states, highlighted with root cell in the monocyte-like cells cluster. B, RNA velocity length D, Proposed hypothetical developmental relationship of pleural CD11b^+^ MNP cells. E, Expression of MNP markers by CD45.1^+^ donor (coloured back for C57BL/6 or red for BALB/c) and CD45.2^+^ (coloured grey) host MΦs from naïve and infected C57BL/6 or BALB/c mice, concatenated from three individual mice. F, Percentage of donor and host CD11b^+^ MNP that are within the SCM population. G, Percentage of donor and host CD11b^+^ MNP that are CD102^+^ (LCM committed). H, Expression of the Converting CM markers CD206, Lyve1 and PDL-2 by host and donor cells. I, Percentage of donor and host CD11b^+^ MNP that are within the terminal LCM population

**Figure S7.**
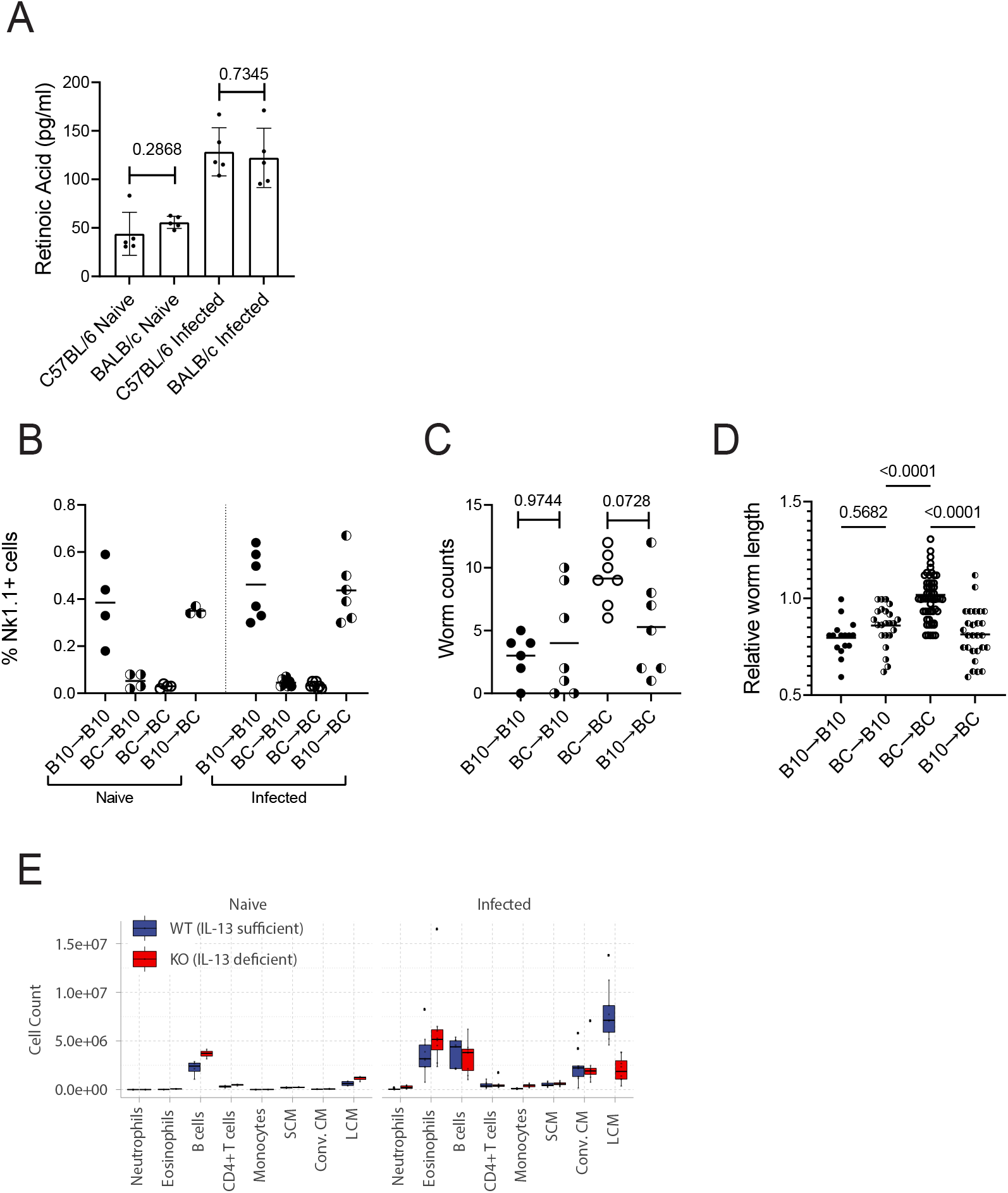
Related to Figure 6 and 7. A, Retinoic acid concentration in the pleural fluid in naïve and infected C57BL/6 and BALB/c mice. B, Expression of the allogenic marker Nk1.1 expressed by NK cells from B10.D2 mice but not BALB/c mice by pleural cavity immune cells following bone marrow transplantation. C, Worm counts at day 34 p.i. from mice that received bone marrow transplantation. p values represent Mann-Whitney U test. D, Relative worm length (normalised for worm sex) at day 34 p.i. from mice that received bone marrow transplantation. p values represent one-way ANOVA with Bonferroni’s multiple comparisons correction. E, Pleural cell numbers by cell type from IL-13 eGFP^-/-^/IL-13 eGFP^-/+^ (IL-13 sufficient) or IL-13 eGFP^+/+^ (IL-13 deficient) mice on day 35 p.i..

## Supplemental information

### Materials and methods

#### Materials table

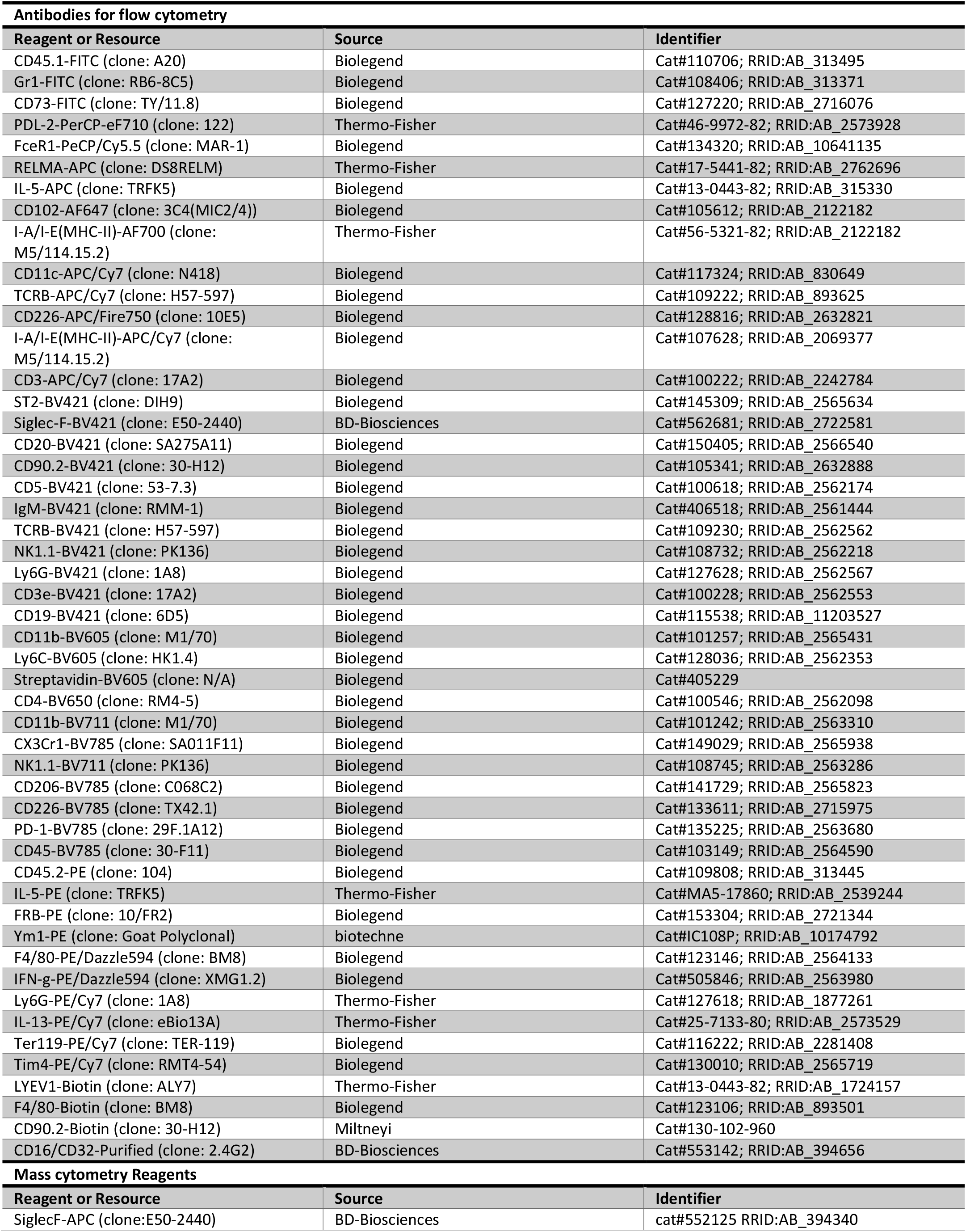
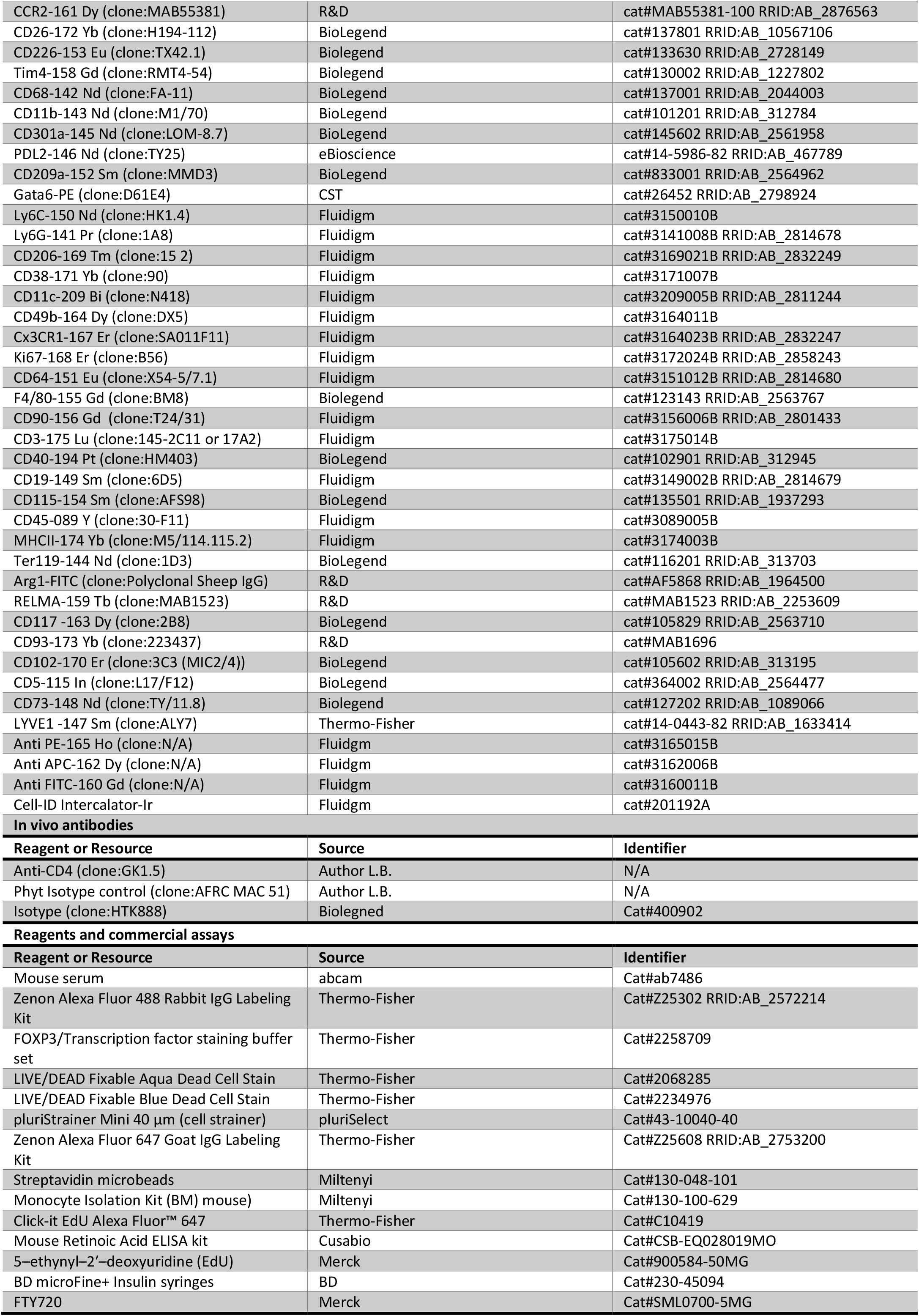
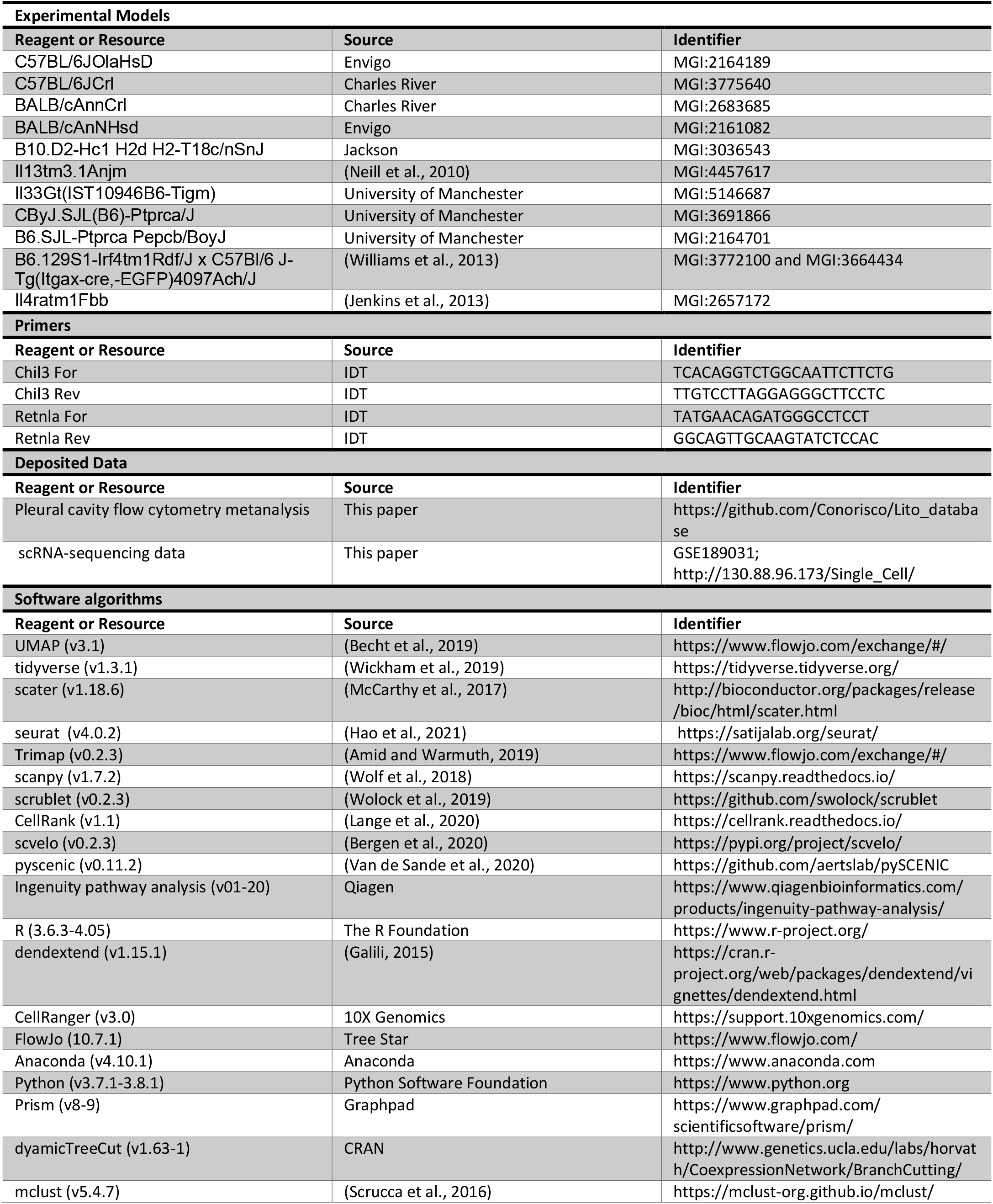

#### Materials availability

Further information and requests for resources and reagents should be directed to the corresponding authors.

#### Materials availability

This study did not generate new unique reagents.

#### Data and code availability

Detailed laboratory notebooks, code and raw data are available from the corresponding author upon request. Links to raw data and analysis pipeline for the collective analysis and scRNA-seq data are available in resource table.

## Methods

### Mouse strains

Female C57BL/6J^OlaHsD^ (*H2*^*b*^) mice and BALB/c^OlaHsd^ (*H2*^*d*^) mice were purchased from Envigo, or bred in-house. C57BL/6J-CR and BALB/c^AnNCrl^, purchased from Charles River were used for a minority of experiments. B10.D2 (B10.D2^*Hc1 H2d H2-T18c*/nSnJ^) which are serum C5 sufficient, carrying the *Hc* locus from C57BL/10Sn mice, were purchased from Jackson laboratories and bred in house. IL-13-eGFP mice (*Il13*^tm3.1Anjm^, backcrossed to C57BL/6J mice) were generated via heterozygous littermate pairings to produce WT, heterozygous and homozygous mice. IL-4Rα ^-/-^ mice (Il4ra^tm1Fbb^, backcrossed to C57BL/6J) were generated by homozygous pairing. IL-33LacZGT (*Il33*^Gt(IST10946B6-Tigm)Girard)^) were generated via heterozygous littermate pairings to produce WT, heterozygous and homozygous mice. CD45.1 BALB/c mice (CByJ.SJL(B6)-^*Ptprc*a^/J) and CD45.1 C57BL/6 mice (B6.SJL^*Ptprca Pepcb*/Boy^J) were generated via homozygous littermate pairings. CD45.1^+^ CD45.2^+^ heterozygous mice were generated via breeding these homozygous strains with C57BL/6J and BALB/c mice. Irf4^fl/fl^; B6.129S1-Irf4^tm1Rdf^/J x C57Bl/6J-Tg(Itgax-cre,-EGFP)4097Ach/J *(Irf4*^*fl/fl*^ *x CD11c-*Cre-GFP) were maintained by heterozygous littermate pairings to produce WT, heterozygous and homozygous mice. Mice were aged between 6 weeks and 11 months of age at the start of each experiment. The age of mice within experiments were matched to within 2 weeks. Female mice made up 83 % of all mice used in the study. Mice were bred in house and maintained in specific pathogen free (SPF) facility. Experiments were in accordance with the United Kingdom Animals (Scientific Procedures) Act of 1986 under the project Licenses 70/8548 and PP4115856. Mice on BALB/c backgrounds which had evidence of spontaneous thymomas were removed from all analysis.

### L. sigmodontis infection

Tropical rat mites, *Ornithonyssus bacoti*, were feed on microfilaria positive Mongolian Jirds (*Meriones unguiculatus*) overnight. Fully engorged mites were then collected and incubated at 27 °C, 75% relative humidity for at least 12 days to allow development to the infective third stage larvae (L3). Mites were then crushed in media (RPMI supplemented with 5% horse serum). 25 L3s were then injected subcutaneously in the scruff of the mouse using a 23G needle in 200 μl of media. Detailed instructions on the maintenance of the life cycle (Fulton et al., 2018) and the immune responses (Finlay and Allen, 2020) *L. sigmodontis* have been published elsewhere.

### In vivo chemical, antibody or cytokine administration

For assessment of cell proliferation mice were injected intraperitoneally (i.p.) with 5–ethynyl–2′– deoxyuridine (EdU) (0.5 mg/mouse) three hours prior to culling of animals. For T cell depletion experiments, 200 μg of anti-CD4 (GK1.5, Rat IgG2b) or isotype control (AFRC, Mac 5.1) in PBS were injected i.p. on day 7 and every 7 days thereafter until the end of the experiment. For FTY720 experiments, FTY720 was administered in the drinking water (final concentration of 2.5 μg/ml) one day before infection. Water pouches were replaced weekly with fresh prepared FTY720 added each time.

### Magnetic Resonance Imaging

To visualise any pleural oedema resulting from infection, post-mortem coronal T2-weighted TurboRARE scans were acquired covering the thoracic cavity. Scans were acquired on a Bruker Avance III console interfaced with an Agilent 7T 16 cm bore magnet using the following acquisition parameters: TR/TE = 5320/33 ms, NEX = 2, RARE factor = 8, echo spacing 11ms, field of view = 30 × 30, matrix size = 512 × 512, voxel size = 0.059 × 0.059 mm^2^, 50 slices, slice thickness = 0.5 mm. To quantify pleural oedema (hyperintense signal), the pixels containing lung and oedema were manually segmented in MRIcron(v1.0) to produce a lung mask. All pixels outside the lung mask were set to zero, and pixels within the masked area set to either 1 or 0 depending on whether they were above or below 2*median of all masked pixel values. Pixels above the threshold (1’s) were classified as oedema and counted. The total oedema volume was calculated for each animal by multiplying the total number of pixels classified as oedema by the pixel volume (0.0017 mm^3^).

### Pleural cell, fluid, and worm isolation

Pleural exudate cells (PLEC) were obtained via washing of the pleural cavity with 2 ml ice-cold PBS followed by 6-8 ml of PBS supplemented with 2 % FCS and 2 mM EDTA using a sterile transfer pipette. Cells were filtered through a sterile pluriStrainer Mini 40 μm cells strainers (pluriSelect) into a 15 ml tube. Worms retained on the filter were removed by back washing with RPMI into a 6 well plate for counting using a dissecting microscope. PBS-only washes were centrifuged, and the supernatant stored in a biobank for future analysis by competitive ELISA for retinoic acid detection using Mouse Retinoic Acid ELISA kit (Cusabio) using manufactures instructions. The cell pellet of the PBS-wash was combined with the cell pellet of the PBS-FCS-EDTA washes and cells were counted using a Nexcelom automated cell counter. For a minority of samples, erythrocytes were lysed prior to cell counting.

### Flow cytometry

Cells were stained in V-bottom 96 well plates in 50 µl reaction volumes kept at 4 °C. Cells were washed by addition of 200 μl of relevant buffer and centrifugated at 300-400g for 5 mins prior to fixation and 700 g for 5 mins after fixation, cells were first washed twice with PBS, and then incubated with anti-CD16/CD32 5 μg/ml (Mouse Fc Block, BD) and LIVE/DEAD Blue or LIVE/DEAD Aqua (ThermoFisher) in PBS for 30 min. Next cells were washed with PBS with 2% FCS and 2 mM EDTA (hereafter FACS buffer) and incubated with antibodies to surface antigens (0.5 µg/ml) with mouse serum (1 in 50). We used a lineage cocktail on the same fluorophore which contained TCR-β, CD20, CD19, Ly6G, Nk1.1, CD90.2, CD5, IgM, CD3ε and Siglec-F, with some experiments using Ly6G on another channel, and some experiments forgoing CD5 and Nk1.1. Optionally, for Intracellular antigen staining, cells were fixed and permeabilized using the Foxp3/Transcription Factor Staining Buffer Set overnight and stained with antibodies against intracellular antigens (1 µg/ml) in Permeabilization buffer (ThermoFisher). For biotinylated antibodies, fluorophore-conjugated streptavidin (Biolegend) was added in the same reaction (1 in 400 from stock). Optionally, for assessment of edU incorporation we used the Alexa Fluor 647-azide using Click-iT Plus EdU Flow Cytometry Assay Kit (ThermoFisher) according to the manufacturer’s instructions. Samples were acquired on LSRFotessa X-20 or LSRFortessa (BD). Exclusion criteria: Samples removed from analysis if they contained staining or acquisition artefacts in key parameters if this made comparison with other samples difficult or if the number of events in the ‘live cell’ gate was less than 1000. Analysis was performed using FlowJo and gating strategies are detailed in Figure S1. Gating strategies and optimal fluorophore and antibody choices were refined through the project and alternative gating strategies exist (Figure S1). To remove staining/acquisition artefacts we used FlowAI Flowjo Plugin (Monaco et al., 2016). Anti-CD115 was not used to identify monocytes/MΦs because in our hands we observed a down regulation of CD115 by MΦs after isolation that was inconsistent between experiments.

### Collective analysis of L. sigmodontis data

Measured variables from 40 independent *L. sigmodontis* infection experiments containing 666 (535 female and 113 male animals with a median age 124 days at end of experiment) mice were compiled into a single dataset containing mice of various backgrounds that were infected or uninfected (Naïve). Genetically modified mice which were homozygous negative (‘wild type’) for the respective mutation were also included for in the analysis. Analysis includes mice from reductionist experiments if these mice were in the control groups, that is they received i.p. injections of PBS, distilled H2O added to drinking water or isotype control antibody. Mice were then filtered to include only those on a BALB/c or C57BL/6 background. An additional 24 mice were removed from analysis for either failing to meet quality control standard of flow cytometric data (i.e. low event count, staining artefacts) or they were deemed to be not infected. We defined a failed infection as those mice which were injected with L3 larva but upon analysis had no live worms, no granulomas, no evidence of moulted worm exoskeleton no increase in PLEC count over baseline and PLEC eosinophils <2% of PLEC. No mice were excluded after day 50 p.i. as parasite clarence rather than failed infection could explain these observations. After data filtering, 360 mice remained (311 female and 49 male, median age 103 days at end of experiment). Metadata included the following categorical data: experiment number, sex, infection status, strain, genetic background, genotype, and supplier. Numerical data included: age, blood score, worm counts, granuloma counts, cell counts, live cell %, L3 infection dose, and numerical outputs from flow cytometric analysis. To ensure consistent inter-experiment flow cytometry data, all experiments retained after filtering were reanalyzed in Flowjo using the gating strategy displayed in Figure S1. Analysis was performed in R using the Tidyverse package libraries. Groups were collated by genetic background irrespective of sex, supplier, and sub-strain. Data visualization was created using the ggplot2 R package. For bar chart summary graphs infected groups included data from day 23 and day 60 p.i. Statistics were performed using the stat_compare_means function from the ggpubr R package. For time course data, naïve samples were given the time of 0. Lines in time course graphs model a polynomial regression with locally estimated scatterplot smoothing (loess), generated using the geom_smooth function from ggplot2.

### Mass Cytometry

PLEC were isolated from naïve and day 36 p.i. *L. sigmodontis-*infected C57BL/6 and BALB/c mice (5 female mice per group). We retained only 3 mice per group and excluded lower quality samples that had blood contamination of the PLEC or absence of worms in infected groups. No buffers stored in glass were used in the protocol. Cells were incubated at 4 °C for all steps in staining volumes of 50 μl in v-bottom plates. Cells were washed in PBS and stained with 2.5uM Cisplatin for 2 minutes and washed with FACS buffer. Next, cells were blocked with mouse serum (1 in 50) and 10 μg/ml anti-CD16/CD32, followed by staining with 3 μg/ml anti-Siglec-F APC. Next following a wash step, metal-conjugate antibodies for surface antigens or APC were added to cells (dilution range 1 in 25 to 1 in 100 from stock). Next, cells were fixed overnight using FoxP3 Transcription Factor fixation/permeabilization buffer followed by following by staining with Arg1-FITC and GATA6-PE together in Permeabilization buffer. After washing with permeabilization buffer, cells were stained with a master mix of metal conjugated antibodies against intracellular antigens, FITC and PE in Permeabilization buffer. Next, cells were washed in permeabilization buffer, and stained with 125 nM cell-ID Intercalator-Ir (Fluidigm) in permeabilization buffer to stain DNA. Finally, cells were washed in FACS buffer and frozen in 10% DMSO plus 90% FBS in a Mr Frosty (Nalgene) at -80 °C. After thawing acquisition occurred on Helios Mass Cytometer (Fluidigm) with EQ Four Element Calibration Bead run normalisation (Fluidgm). Metal conjugation of purified antibodies not purchased from Fluidigm was performed by the University of Manchester Flow Cytometry Core facility. Analysis was performed in Flowjo Firstly, EQ Four Element Calibration Beads were removed for dual staining of 165Ho and 140Ce. Next, viable immune cells were gated for positivity for CD45 (Y89) and negativity for Cisplatin (195Pt). Doublets were removed as follows: firstly, high DNA signal cells were removed (191Ir/191Ir); secondly, cells with high signal for the Event Length, Centre, Residual or Width parameters were removed. Next, all unused parameters were removed using the R package Premessa. Next cells were down sampled to 7916 cells per sample using Downsample Flowjo plugin and merged using Flowjo Concatenate function. Parameters were manually rescaled in Flowjo using an arcsine transformation. UMAP was generated using all parameters corresponding to antibodies using the UMAP Flowjo Plugin (nearest neighbours=15). Cell types were gated using sequential gates/NOT gates in the following order: T cells positive for CD3 (175Di), B cell positive for CD19 (149Sm), Eosinophils positive for Siglec-F (162Dy) and CD11b (143Nd), Neutrophils positive for Ly6G (141Pr) and CD11b (143Nd), Mast cells positive for CD117 (163Dy), ‘Other lymphoid’ positive for CD90 (156Gd), ‘Non-MNP DC’ positive for CD11c (209Bi) and I-A/I-E (174Yb) but negative for CD11b (143Nd) or Ly6C (150Nd), ‘Myeloid’ positive for CD11b (143Nd), Monocytes, positive for Ly6C (150Nd), LCM positive for F4/80 (155Gd) and CD73 (148Nd)(with further discrimination by Tim4 expression), Converting CM positive for Lyve1 (147Sm), the remaining cells were classified as SCM. Expression values displayed on UMAP embeddings were generated using Flowjo heatmap statistic feature based on manual arcsine transformed expression values.

### Single cell RNA-sequencing

For the input of the scRNA sequencing, we chose the day 35 p.i. timepoint as this is when worm killing begins in C57BL/6 mice and monocyte influx begins in infected BALB/c mice (Finlay and Allen, 2020). The total PLEC of 3 naïve C57BL/6 mice and 3 naïve BALB/c mice were pooled prior to sorting to maximise yield and minimize group variation. For infected groups cells were pooled equally following cell sort. For the C57BL/6 Infected group, the PLEC from 4 mice were pooled. For the BALB/c Infected group, the PLEC of 3 mice were pooled with 2 additional mice excluded due to blood contamination of the lavage. PLEC cells were strained through a 40 μm filter and stained with Live/Dead and fluorophore-conjugated antibodies against CD11b, CD45.2, CD11b, Ly6C, F4/80, I-A/I-E and Ter-119 and PDL-2 along with a lineage cocktail (TCR-β, CD20, CD19, Ly6G, Nk1.1, CD90.2, CD5, IgM, CD3ε and Siglec-F). Live single, lineage^-^CD45.2^+^Ter^-^119^-^CD11b^+^ cells were sorted into Eppendorf’s containing PBS with 2 % FBS. Cells were washed in PBS, counted and 15700 cells per sample were loaded onto a 10X Genomics Chromium controller following manufacture’s protocol to create Gel-Bead in Emulsions (GEM). Cell viability prior to loading was > 98%. 4 Separate cDNA libraries were created using the Single Cell 3’ Library Version 2 Kit. After final preparation and QC performed, libraries were pooled to equal molarity. Library preparation was performed by University of Manchester Genomics technology core facility. The pooled Library was sequenced with paired-end reads using a full S2 lane of a NovaSeq-6000 (Illumina) by Edinburgh Genomics yielding 1750 million reads.

### scRNA-seq data processing

Sequencing reads in fastq format were aligned to the mm10 mouse transcriptome using Cell Ranger (10X Genomics) count function, quantifying Unique Molecular Identifiers (UMI) for each gene associated with each cell barcode. Samples were integrated and normalised using the Cell Ranger aggr function producing a gene versus cell expression matrix. Downstream analysis was performed in R using the Bioconductor suite of single cell analysis tools scater, scran and singlecellexperiment. The merged counts were first normalised by library size and CPM calculated using the calculateCPM function along with a log2 transformation with an offset of 1, hereafter ‘log-normalization’. Poor quality cells were filtered by removing those with a low feature count (3*Median absolute deviations (MAD)), low number of genes (3*MAD) and percentage mitochondrial reads (7*MAD, equating to 9% mitochondrial reads). We chose the higher cut-off for mitochondrial genes as cells from the C57BL/6 infected group had a higher proportion of reads mapping to mitochondrial reads fitting with previous reports that M(IL-4) signalling leads to mitochondrial biogenesis (Odegaard et al., 2007). Genes were filtered to retain those with an average count of 0.005 per cell. Filtering resulted in 10,522 genes and 25,904 cells being retained for downstream analysis. Cells were renormalised after filtering.

### SCENIC Transcription factor analysis and dimension reduction

A single cell transcription factor (TF) gene regulatory network was inferred using PySCENIC (Van de Sande et al., 2020), the python implementation of SCENIC (Aibar et al., 2017), using log-normalized counts described above. First, weighted TF-gene co-expression modules were inferred using the regression-per-target GRNBoost2 algorithm (https://github.com/tmoerman/arboreto). Next, indirect target genes were pruned to retain those target genes which contain matching transcription factor DNA binding motif within their regulatory regions. This was achieved using all the mouse mm9 version 9 (mc9nr) cisTARGET databases (https://resources.aertslab.org/cistarget/). Finally, regulon activity was quantified by calculating an enrichment score for the target genes within each regulon using AUCell. The pipeline generated 251 regulons, of these 56 were further removed for having low activity in our dataset.

### Dimension reduction and clustering

SCENIC regulon AUC scores were used for dimension reduction and clustering. The 195 regulon scores were scaled and used to generate an UMAP embedding with the arguments: nearest neighbours = 100 and minimum distance = 0.15 using the umap R package without an intermediate PCA step. Scaled regulon AUC scores were clustered using Ward’s method of hierarchal clustering of squared Euclidean distance using the hclust R package (implementation ‘Ward.D2’). To generate metaclusters we used the cutreeDynamic wrapper function from the R package dynamicTreeCut using the default cutoff of 99% of the maximum joining hights of the dendrogram. This resulted in 9 metacluters which were manually annotated based upon the top relative high expressed genes in each cluster. One cluster with high transcripts for genes typically expressed by B cells was removed from downstream analysis. Dendrogram visualisation was created using the dendextend R package. Data visualisations for scRNA-seq data was made using the ggplot2 R package.

### Differential gene expression, pathway analysis and scoring of biological processes

Differential gene expression analysis was performed in using the Seurat R package following conversion of the singlecellexperiment object to a SeuratObject. This included a data normalisation that differed from that calculated using the Bioconductor pipeline. Seurat’s logNormalize function divided raw gene counts in each cell by total counts in that cell and multiplied this by a scale factor of 10000 and then natural log transform with an offset of 1. Seurat normalised counts were not batch corrected. Cluster-specific differential gene expression was calculated using FindMarkers or FindAllMarkers functions using the arguments minimum percentage expression = 0.25 and log fold change threshold = 0.25. For differential regulon activity FindMarkers or FindAllMarkers functions were used without minimum thresholds. Genes which passed these thresholds were used as input to Ingenuity Pathway Analysis, using log fold change as the analysis metric. IPA was run using standard settings and all experimentally confirmed mouse reference datasets (without cell line datasets) were included. Scoring of biological processes was performed as described in (Xie et al., 2020). Scores were the mean log-normalized gene expression of all genes in a given gene list. Bi-directional scores were generated as the weighted average of z-scores of genes in a gene list where each gene given a -1 or 1 to mirror downregulated or upregulated genes in that gene list. Datasets used to generate Immgen-likeness scores were downloaded from <u>Immunological Genome Project (immgen.org)</u> (version 1 microarray) originally generated by the Randolf laboratory (Gautiar et al., 2012). The specific datasets we used were ‘MF.II-480hi.PC’ for LPM and ‘MF.II+480lo.PC’ for SCM. From this we took the top 100 differentially expressed genes in each direction to generate weighted scores. IL-4c scores were generated from a gene list of the top 50 genes upregulated in peritoneal LCM following *in vivo* administration of IL-4c (Gundra et al., 2014). Peritoneal MΦ-specific gene (PMSG) and their GATA-6 dependency gene lists were taken from (Okabe and Medzhitov, 2014). Type 1 interferon scores were generated using the genes within the Gene Ontology term GO:0071357 filtering for mouse only.

### RNA velocity and CellRank fate mapping analysis

Trajectory analysis and cell fate inference was performed using RNA velocity (La Manno et al., 2018). using the python packages velocyto, scvelo (Bergen et al., 2020) and CellRank (Lange et al., 2020). RNA velocity calculates the ratio of spliced to unspliced mRNAs for each gene. Velocities are vectors in a given gene expression space that demonstrate the directionality and speed (velocity ‘length’) for each cell with positive velocities in a cell showing that that gene is being upregulated by having higher unspliced to spliced ratios (relative to steady state), and opposite true for genes being downregulated. A vector field constructed from all velocities can be projected in low dimensional space to determine the directional flow of cell states. CellRank adapts this data to predict the cell fate of each cell from an initial state to terminal states. Calculation of spliced and unspliced reads was produced using the velocyto.py command line tools using the original fastq sequencing files for each sample independently. The resulting loom files of spliced-unspliced gene counts were merged and cell barcodes were loaded into python using the scvelo package, creating an anndata object (Bergen et al., 2020). Next, we filtered the dataset to reflect the bioconductor analysis pipeline and imported the SCENIC regulon based UMAP embedding from R. The B cell cluster was removed from all further analysis. C57BL/6-like LCM and BALB/c-like LCM were combined to create a ‘Naïve LCM’ cluster. For analysis of naïve samples, only the C57BL/6 sample was used and the proliferating, M(IL-4) and Converting CM clusters were removed. We next ran the datasets through the scanpy python package pipeline using default parameters which including a normalisation step identical to that used by Seurat. Highly variable genes were identified in scanpy and scanpy regress_out function was used to regress out total counts per cell. PCA was calculated by scanpy with default parameters and the top 30 PC were used to calculate a neighbourhood graph with nearest neighbours=30. As doublets might be incorrectly identified as transitionary cell states we computationally removed doublets using the scrublet python package (Wolock et al., 2019). Next, the anndata object was passed to scvelo python package (Bergen et al., 2020). This first ran the filter_and_normalize function with the following arguments min_shared_counts=30, n_top_genes=2000. We also a supplied a list of genes to retain that were of biological interest to us: *Lyve1, Icam2, Cd74, Pdcd1lg2, Adgre1, Itgam, Itgax, Ccr2, Dpp4, Nt5e, Retnla, Chil3, Arg1, Folr2, Timd4, Marco, Mertk, Ly6c1, Ly6c2, Dpp4, Selp, F5, F10, Gata6, Bhlhe40, Cebpb, Cxcl13, Pf4, Apoe, Cd36, Mgl2, Cd209a, Cd209d, Prg4, Saa3, Alox15, and Napsa*. This was followed by the scvelo moments function and then the core scvelo velocity and velocity_graph functions using the standard stochastic mode, thus creating a matrix of RNA velocities for each cell. The velocity_graph function was used to project the combination of these velocities in lower dimensional space, and velocity_embedding_stream to project this on a PCA or UMAP embedding. We used a cut-off of 3.5 to display only higher velocities on the embedding plots. For the final part of trajectory analysis we used CellRank (Lange et al., 2020) for inference on initial and terminal cell fates. Terminal states were identified using the terminal_states using the regulon clusters as the cluster key, where this resulted in multiple terminal states in the same cluster n_states argument was set to 2. Next the initial_states function identified the initial states and fate maps computed by the lineages function. To plot cluster aggregated cellular fates we used the cluster_fates function to produce absorption possibilities or the likelihood that cells in that cluster will progress to a given terminal state. To visualise directed cluster connectivity we used CellRank recover_latent_time and paga functions to create a directed partition-based graph abstraction (PAGA) graph (Wolf et al., 2019) that was projected on PCA space.

### Intrapleural injections

Syringes (BD microFine+ Insulin syringes) were prepared in advance by drawing 100 μl of PBS cell suspension into a 27 g low dead volume insulin syringe and air expelled. The final 5 mm of the needle tip was crooked at an 80° angle using a sterile forceps with bevel pointing out. The mouse was anaesthetised with isoflurane and the left thoracic region shaved with an electric clipper and sterilised. Injections were performed by piercing the skin at an intercostal space of lower left sternum with the point of the needle and injecting the solution, with the crook limiting the penetration to 5 mm. Mice were observed for pneumothorax for 30 mins after procedure. Mice that received improper injections (e.g. intradermal as evidenced by skin bulge at injection site) were removed from analysis.

### Monocyte transfers

Bone marrow was isolated from the tibia/femur bones of male Pep3 C57BL/6 CD45.1^+^ and BALB/c CD45.1^+^ and single cell suspension prepared after lysis of red blood cells. These cells were stained for FACS with Live/Dead and fluorophore conjugated antibodies against CD45.1, Ly6C, CD45.2 and a lineage cocktail containing antibodies (Ter119, FcεR1, CD3ε, TCR-β, B220, Nk1.1, Siglec-F, Ly6G). Single, live, CD45.1^+^, lineage^-^CD11b^+^Ly6C^high^ mature monocytes were sorted on a FACSAria Fusion (BD). 8.5 ×10^4^ sorted monocytes were injected intrapleurally into male naïve and day 15 p.i. CD45.2^+^ BALB/c and CD57BL/6 in a strain matched manner. PLEC were isolated on day 35 p.i. or 21 days post transfer and were analysed by flow cytometry for expression of CD45.1, PDL-2, CD102, I-A/-I-E, CD11c, Ly6C, Lyve1, CD11b, CD206, CD45.2, F4/80, Tim4 and a lineage cocktail (Siglec-F, TCR-β, Ly6G and CD19). Transferred cells were identified as CD45.1^+^Lineage^-^CD11b^+^ cells and donor CD11b^+^ MNPs as CD45.1^-^CD45.2^+^Lineage^-^CD11b^+^ cells. For experiments using 1:1 dual transfer of C57BL/6 and BALB/c monocytes into the same CD45.2^+^ host, bone marrow monocytes were instead sorted from bone marrow using a bone marrow monocyte Isolation kit (Miltenyi). Additionally, to discriminate the transferred cells from in a strain-specific manner, donor BALB/c CD45.1^+^CD45.2^+^ heterozygous and C57BL/6 CD45.1^+^ homozygous were used. PLEC cells were analysed after 4 days before onset of tissue rejection.

### Bone marrow transplantation

Female BALB/c and B10.D2 mice, which share the H^2d^ MHC haplotype were used to create strain-matched and strain mismatched bone marrow chimeras. To prevent infection, 2 days before irradiation and until 25 days post irradiation mice were placed on 0.3 % Enrofloxacin (Baytril 10% Oral, Bayer) in acidified drinking water and were supplemented with irradiated diet as mash. On Day 0, BALB/c mice were administered a total of 925 Gy, while B10.D2 were given a total of 1100 Gy, given the higher susceptibility of BALB/c mice to irradiation. These doses were given as 2 doses 2 hours apart using an X-ray source (Xstrahl RS320, provided by Epistem Ltd). 6 hours later, mice were injected with 3.25 million live bone marrow cells from naïve BALB/c or B10.D2 donors in PBS by i.v. injection. To remove potentially allograft reactive T cells, bone marrow was first depleted of CD3 and CD90.2 positive cells by negative magnetic separation, using CD3-biotin, CD90.2-biotin, streptavidin microbeads and LS columns. Mice were weighed daily for first 7 days and 3 times per week thereafter. Mice that lost 20% of initial weight were culled. Of a total of 48 mice, 2 B10.D2 and 5 BALB/c mice were removed from the experiment. This resulted in the final group sizes of B10.D2 into B10D2 of 4 naïve and 6 infected, BALB/c into B10.D2 of 4 naïve and 8 infected, BALB/c into BALB/c of 4 naïve and 7 infected and B10.D2 into BALB/c of 3 naïve and 6 infected. On day 42 post-irradiation mice were infected with *L. sigmodontis*. Mice were culled on day 77 post irradiation which was day 34 p.i.. Chimerism was determined by expression of the B10.D2 alloantigen Nk1.1 expression by NK cells in the blood and pleural space.

## Notes

https://github.com/Conorisco/Lito_database

https://www.ncbi.nlm.nih.gov/geo/query/acc.cgi?acc=GSE189031

http://130.88.96.173/Single_Cell/

### Competing Interest Statement

The authors have declared no competing interest.

